# Functional Characterization of Target of Rapamycin (TOR) Signalling in Physcomitrella

**DOI:** 10.1101/2025.11.04.686496

**Authors:** Elie Saliba, Sebastian N. W. Hoernstein, Nico van Gessel, Alexander Sentimenti, Karoline M. V. Höß, Juliana Parsons, Eva L. Decker, Pitter F. Huesgen, Henrik Toft Simonsen, Ralf Reski

## Abstract

Target of rapamycin (TOR) is a conserved protein kinase and an important signalling hub in eukaryotes. The moss Physcomitrella (*Physcomitrium patens*) is a model organism for plant physiology, development, and evolution. However, little is known about TOR signalling in non-vascular plants, including Physcomitrella. Here, we report the effects of inhibiting TOR signalling in Physcomitrella. We identified and characterized Physcomitrella 12-kD FK506-binding protein (FKBP12), which binds TOR in the presence of rapamycin. Whereas the growth of wild-type protonema is unaffected by rapamycin, overexpressing endogenous *FKBP12* rendered the plant susceptible to the inhibitor in a dose-dependent manner. Likewise, protonema growth was inhibited when the TOR-specific inhibitor AZD8055 was present in the culture. We show that rapamycin and AZD8055 have pleiotropic effects, as they delay cell cycle progression and development, induce chlorosis, inhibit photosynthesis, and alter total protein content. Additionally, we identified and characterized PpTOR, PpLST8, and PpRAPTOR, key components of TOR complex 1 (TORC1), whereas RICTOR and mSIN1 of TORC2 are not encoded by the *P. patens* genome. We found that PpLST8 substitutes its homolog in yeast to allow cell growth. Physcomitrella mutants were generated with a conditional downregulation of *PpTOR*, *PpLST8*, and *PpRAPTOR*, respectively. They were impaired in growth. Finally, we show TOR-dependent phosphorylation of a well-known TOR phospho-target, the ribosomal protein RPS6, via LC-MS/MS. Collectively, our results show that Physcomitrella growth and development is positively controlled by a conserved TOR kinase. We suggest to further dissect TOR signalling in Physcomitrella in order to elucidate signalling integration via TORC1 in plants.

**Key message:** The evolutionary conserved TOR kinase positively controls growth and development of the moss Physcomitrella, development and function of its chloroplasts, its protein synthesis and cell cycle progression.

## Introduction

All living organisms rely on sensing and responding to their surroundings. The information that is taken up by a significant number of external and internal sensors has to be processed and integrated. Depending on the nature of the information, this is done by complex macroscopic structures (e.g., the trigger hairs on bladderworts or the eyes of animals) or molecular sensors acting through signalling pathways within the cell. One of the most crucial and conserved signalling hubs in eukaryotes is Target of Rapamycin (TOR), best known for its role in controlling metabolism, cell growth, proliferation and survival, mainly in response to nutrients, energy, and environmental stresses (Loewith and Hall 2011; Liu and Xiong 2022; Wang et al. 2025).

TOR is an atypical serine/threonine kinase and a member of the phosphatidylinositol (PI) 3-kinase-related protein kinases (PIKKs) family (Keith and Schreiber 1995). It was originally identified in a genetic screen for yeast (*Saccharomyces cerevisiae*) mutants that could grow in the presence of rapamycin (Heitman et al. 1991a), a macrolide antibiotic from the bacterium *Streptomyces hygroscopicus* (Sehgal et al. 1975; Vezina et al. 1975). TOR proteins are found in most eukaryotes and exhibit high amino acid (aa) sequence similarity (Maegawa et al. 2015; Tatebe and Shiozaki 2017; Liu et al. 2025). They contain five functional domains: HEAT (Huntington, Elongation factor 3, regulatory subunit A of PP2A, TOR1) repeats at the N-terminus, FAT (FRAP-ATM-TRRAP) and FRB (FKBP-Rapamycin-Binding) in the middle of the protein, and the kinase and FATC (Carboxy-terminal FAT) domains at the C-terminus (Tatebe and Shiozaki 2017). Rapamycin inhibits TOR by forming a ternary complex with the highly conserved 12-kDa FK506-binding protein (FKBP12) and the FRB domain, thus restricting its kinase domain (Heitman et al. 1991b; Choi et al. 1996; Yang et al. 2013).

In yeast and mammals, TOR is the catalytic subunit of two high-molecular-mass complexes, TORC1 and TORC2, with distinct architecture and function (Loewith et al. 2002). Yeast TORC1 is formed by either TOR1 or TOR2, LST8 (Lethal with Sec Thirteen 8), KOG1 (Kontroller of Growth 1), and TCO89, while TORC2 exclusively contains TOR2, LST8, AVO1/2/3 (Adheres Voraciously (to TOR2) 1/2/3), and BIT2/61 (Binding Partner of TOR2 protein) (Loewith et al. 2002; Reinke et al. 2004). Mammalian mTORC1 includes mTOR, mLST8, RAPTOR (Regulatory-Associated Protein of TOR), the homolog of yeast KOG1, and two additional proteins, DEPTOR (DEP domain-containing TOR-interacting protein) and PRAS40 (Proline Rich AKT Substrate 40 kDa). Similar to mTORC1, mTORC2 shares core components (mTOR, mLST8, and DEPTOR) and includes the yeast AVO1 and AVO3 homologs mSIN1 and RICTOR, along with PROTOR1/2, equivalents of BIT2/61 (Loewith and Hall 2011; Liu and Xiong 2022). TORC1, but not TORC2, is sensitive to rapamycin (Loewith et al. 2002). This feature was instrumental in elucidating the function of these complexes, with TORC1 regulating transcription, translation, metabolism, and autophagy to ensure cell growth and survival, while TORC2 has specific roles in actin cytoskeleton organization and cell wall integrity (Loewith et al. 2002; Loewith and Hall 2011).

In algae and flowering plants, the composition of TOR complexes is less well understood, with no evidence for the existence of TORC2 (Liu et al. 2025). The plant TOR complex consists of TOR, LST8, and RAPTOR. In Arabidopsis (*Arabidopsis thaliana*), which has one *TOR*, two *RAPTOR*, and two *LST8*, with only *RAPTOR1* and *LST8-1* being predominantly expressed, homozygous *Attor* null mutants are embryo lethal (Menand et al. 2002), whereas *Atraptor1* or *Atlst8-1* knock-out plants show embryo developmental arrest and severe growth defects, respectively (Deprost et al. 2005; Moreau et al. 2012). At the cellular level, studies mainly conducted in Arabidopsis revealed that TOR responds to nutrients, energy, light, hormones, biotic and abiotic stresses to regulate transcription, translation, metabolism, autophagy, and hormone signalling (Liu et al. 2025; Wang et al. 2025). At the organismal level, it affects all Arabidopsis developmental stages (Liu et al. 2025). These studies, along with others carried out in algae, have been crucial in understanding the functional evolution of TOR. They also uncovered plant-specific regulations and functions, such as TOR activation by light and auxin through the plant-specific ROP2 GTPase (Moreau et al. 2012; Li et al 2017), and its critical role in promoting photosynthetic activity (Couso et al. 2021; D’Alessandro et al. 2024).

In all examined species, TOR mediates its function by fine tuning the activity via phosphorylation of a large array of downstream effector proteins (Battaglioni et al. 2022; Liu et al. 2025). One key effector that is primarily activated by TORC1 is p70 ribosomal S6 kinase 1 (S6K1), which further phosphorylates its primary substrate ribosomal protein S6 (RPS6) (Jacinto and Lorberg 2008; Battaglioni et al. 2022; Liu et al. 2025). This TORC1-S6K1-RPS6 signalling branch mainly regulates ribosome biogenesis and protein translation (Jacinto and Lorberg 2008; Battaglioni et al. 2022; Liu et al. 2025). In humans, S6K1 is predominantly phosphorylated by mTORC1 on threonine 389 (Kim et al. 2002), while RPS6 mostly undergoes phosphorylation on serines 240 and 244 in an mTORC1-S6K1-dependent manner (Roux et al. 2007). In Arabidopsis, S6K1 and RPS6, along with their TOR-dependent phosphorylation sites, are conserved. AtS6K1 and AtS6K2 are phosphorylated on threonines 449 and 455, respectively (Schepetilnikov et al. 2011). AtRPS6A and AtRPS6B are phosphorylated on Ser240, equivalent to the Ser240 in HsRPS6, and have also been found phosphorylated on Ser237 (Dobrenel et al. 2016). Although these findings underscore the deep evolutionary conservation of TOR signalling mechanisms, our understanding of this pathway in non-vascular land plants, such as the moss Physcomitrella (new species name *Physcomitrium patens*), remains less well explored.

Physcomitrella is a rather small moss in the Funariaceae family (Ostendorf et al. 2021), and a prime model system for studying the biology of non-vascular plants, evolutionary developmental (evo-devo) biology, and cell biology (Reski 2018; Rensing et al. 2020). This prominence is due to its simple body plan (Reski 1998), ease of genetic manipulation due to a yeast-like high rate of homologous recombination in mitotic cells (Strepp et al. 1998; Hohe et al. 2004; Ermert et al. 2019), and its evolutionary position (Beike et al. 2015). One remarkable feature of Physcomitrella is its ability to grow in the form of a filamentous tissue, the protonema, in suspension cultures (Hohe and Reski 2005). This system offers a homogenous and rapidly proliferating biological platform that is also amenable to large-scale propagation under controlled conditions, providing a uniform population of cells (Hohe et al. 2002) for experimental manipulation, genetic engineering, and physiological studies (Decker and Reski 2007). Moreover, the addition of chemicals, such as hormones or inhibitors, leads to a uniform exposure of every single cell without concentration gradients or effects of a gellant (Reski and Abel 1985; Hadeler et al. 1995). Consequently, Physcomitrella protonema in suspension is ideal for molecular genetics, cell biology, and high-throughput functional analyses (Egener et al. 2002; Schween et al. 2005). By capitalizing on these features, Physcomitrella provided important new insights into the evolution of developmental processes (Khraiwesh et al. 2010; Horst et al. 2016; Resemann et al. 2021; Kozgunova et al. 2022; Hoernstein et al. 2023; Maekawa et al. 2025). Therefore, dissecting TOR in Physcomitrella will contribute to clarifying how this signalling hub operates across the green lineage, informing comparative studies between algae and land plants.

This study reports TOR signalling inhibition in the moss Physcomitrella, thus addressing a major gap in our understanding of TOR signalling evolution and function in non-vascular plants. We identified and characterized core components of this pathway, including FKBP12, TOR, LST8, RAPTOR, and RPS6, and demonstrated their evolutionary conservation. Specifically, we established a Physcomitrella genetic system sensitive to rapamycin, determined optimal conditions for inhibitor sensitivity experiments using rapamycin and AZD8055, an ATP-competitive inhibitor of mTOR (Chresta et al. 2010), constructed conditional RNAi Pptorc1 mutants, and found PpTOR-dependent phosphorylation of PpRPS6 at Ser241 as an essential step toward developing a TOR kinase activity assay.

## Results

### Physcomitrella growth is insensitive to rapamycin

As TOR is a master regulator of eukaryotic cell and organismal growth (Liu et al. 2025), we investigated whether its role in growth promotion is conserved in this moss. To start with, we added rapamycin to protonema on solid medium or in a suspension culture, and evaluated plant growth. We found that Physcomitrella growth is insensitive to rapamycin. Neither the addition of 10 µM of the inhibitor, which is 100-fold higher than the one used to inhibit yeast growth (Heitman et al. 1991a), nor the treatment for a long period of up to 18 days impacted biomass accumulation (**Figure 1A**). Likewise, Physcomitrella growth was not affected by rapamycin in suspension cultures (**Figure 1B**). The fact that Physcomitrella is insensitive to rapamycin has been stated before (Menand et al. 2002), however, without disclosing culture conditions or reporting data.

**Fig. 1.**
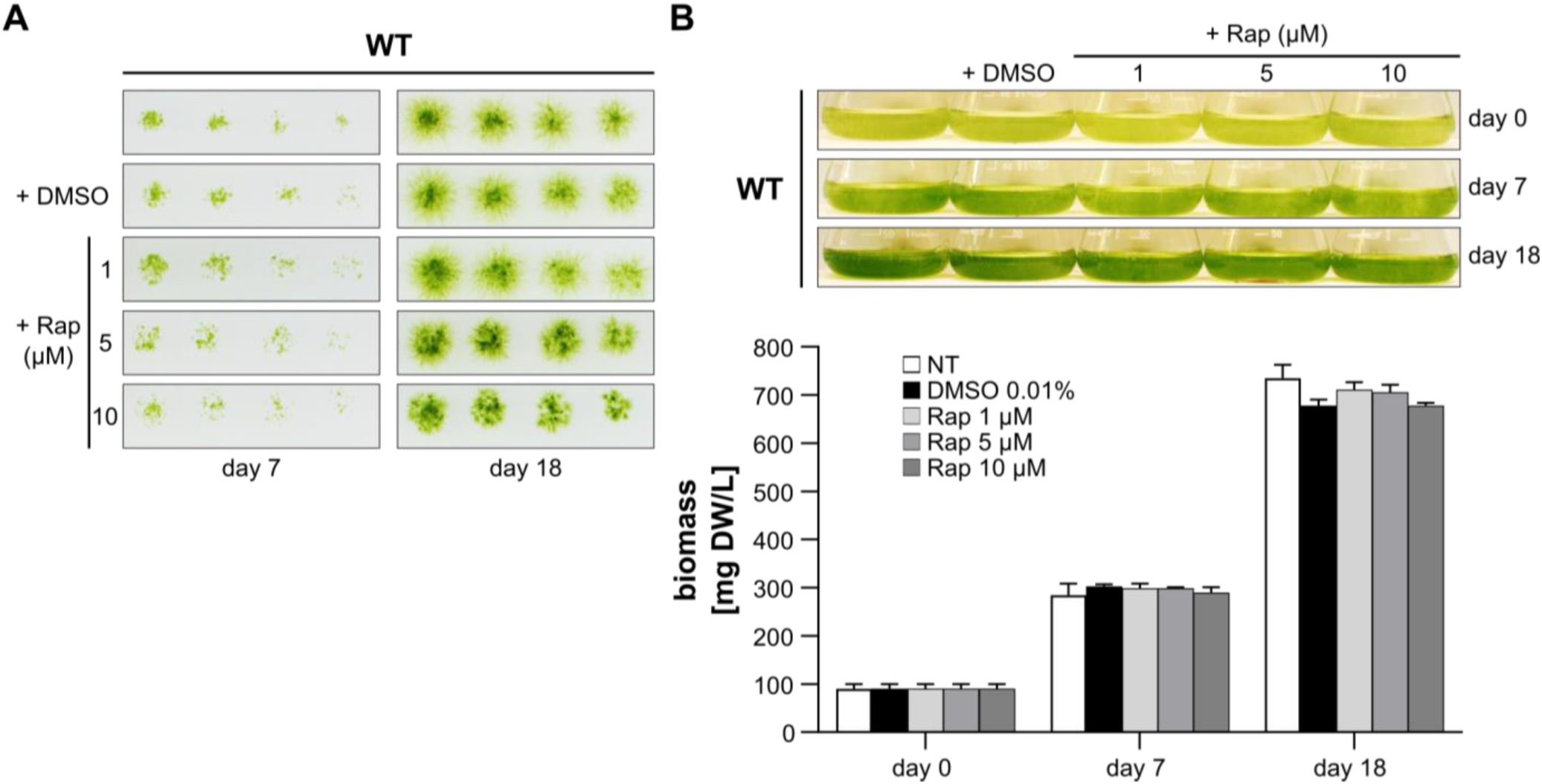
Physcomitrella growth is insensitive to rapamycin. **A** Wild-type protonema from a weekly sub-cultured suspension was prepared at a density of 400 mg DW/L in Knop ME pH 5.8. Serial two-fold dilutions were conducted and 15 µL were spotted on solid Knop ME pH 5.8 plates supplemented with medium (NT=not treated), DMSO 0.01%, or rapamycin (Rap) 1, 5, or 10 µM. Plates were photographed at day 7 and 18 of incubation under 70 µmol photons m^-2^ s^-1^ at 22 °C in a 16 h/8 h light/dark cycle. **B** Same protonema as in panel **A**, except that they were prepared at a density of 100 mg DW/L. Aliquots of 25 mL were established in 100 mL Erlenmeyer flasks and subjected to the same treatments as in panel **A**. Flasks were incubated under 70 µmol photons m^-2^ s^-1^ at 22 °C in a 16 h/8 h light/dark cycle with shaking at 125 rpm. Top: Flasks were photographed at day 0 and days 7 and 18 following incubation. Bottom: Plant material from flasks in panel above were collected and biomass was determined. Bars represent averages of three biological replicates (n=3); error bars represent standard deviations (SD).

Insensitivity to rapamycin appears to be widespread in algae and vascular plants. For instance, growth of Arabidopsis seedlings on solid medium is insensitive to 10 µM rapamycin (Sormani et al. 2007; Ren et al. 2012). Similarly, growth of broad bean (*Vicia faba*) and rice (*Oryza sativa*) is not inhibited, and only high concentrations of the inhibitor could moderately retard growth of tomato (*Solanum lycopersicum*) seedlings (Xu et al. 1998; Xiong et al. 2016; Bakshi et al. 2017). Two algae, the red alga *Cyanidioschyzon merolae* and the green alga *Auxenochlorella pyrenoidosa*, are also insensitive to rapamycin (Imamura et al. 2013; Zhu et al. 2022). Several reasons were suggested: 1) inability of rapamycin to enter plant cells, 2) inability of the FKBP12-rapamycin complex to bind the FRB domain in plant TOR, and/or 3) inability of plant FKBP12 to bind rapamycin or TOR due to a reduced affinity to the drug or a low *FKBP12* expression level. Despite the fact that Arabidopsis seedlings are slightly sensitive to rapamycin when submerged in a liquid medium (Xiong and Sheen 2012), it is unlikely that the insensitivity of algae and plants to the inhibitor is due to poor uptake, or a default in FKBP12-rapamycin-FRB complex formation. In line with this, growth of the green alga Chlamydomonas is sensitive to rapamycin (Crespo et al. 2005). Furthermore, the FRB domain of AtTOR was able to form a complex with rapamycin and ScFKBP12 (Sormani et al. 2007). Moreover, Arabidopsis seedlings became moderately sensitive to rapamycin when *AtFKBP12* expression was increased, a phenotype that is enhanced when human, yeast, or tomato *FKBP12* were ectopically overexpressed (Ren et al. 2012; Xiong and Sheen 2012). In line with this, Chlamydomonas was more sensitive when aa residues in the rapamycin-binding pocket of CrFKBP12, namely glycine 53 or glutamate 54, where substituted with their human or yeast equivalents (Crespo et al. 2005). This suggests that a reduced affinity of plant FKBP12 to rapamycin and/or a low expression level of endogenous FKBP12 are behind plant insensitivity to rapamycin.

### FKBP12 is conserved in Physcomitrella and functional in yeast

A BLASTP search with AtFKBP12 as a query identified Physcomitrella FKBP12 with 76% sequence identity (**Figure 2A**). It is encoded by a single gene, *Pp3c1_27870*, on chromosome 1. A query of the *PEATmoss* gene expression database (https://peatmoss.plantcode.cup.uni-freiburg.de; Fernandez-Pozo 2023) revealed that it is lowly expressed in all tissues (**Supplementary Figure S1**). To evaluate if Physcomitrella’s insensitivity to rapamycin is due to a reduced affinity of PpFKBP12 toward the inhibitor, and/or low expression, we first analysed whether aa residues involved in the interaction between HsFKBP12 and rapamycin are conserved in PpFKBP12. Four of these amino acids (Tyr27, Asp38, Gln53, Glu54) are not present in PpFKBP12 (**Figure 2A**, black arrows). Instead, a cysteine, a tryptophan, a glycine, and an arginine occupy these positions, respectively. Hence, it is possible that the presence of these aa impairs the interaction between PpFKBP12 and rapamycin in Physcomitrella. To test this hypothesis, we generated five PpFKBP12 mutants with single or multiple substitutions of these aa with their human counterparts (**Figure 2B**; mutants PpFKBP12^C26Y^, PpFKBP12D^W40D^, PpFKBP12^C26Y-W40D^, PpFKBP12 ^M56K-G57Q-R58E^, and PpFKBP12^C26Y-W40D-M56K-G57Q-R58E^). As for Gly57 in PpFKBP12, the two aa on both sides were additionally replaced by their human homologs (PpFKBP12^M56K-G57Q-R58E^ and PpFKBP12^C26Y-W40D-M56K-G57Q-R58E^ variants). This decision was adopted since a CrFKBP12 mutant bearing a lysine, a glutamine, and a glutamate at the respective positions conferred more sensitivity of Chlamydomonas cells to rapamycin than the WT protein (Crespo et al. 2005). Two additional mutants were generated, in which Arg58 was not substituted with its human homolog, but rather with its yeast one (**Figure 2B**; PpFKBP12^M56V-R58Q^ and PpFKBP12^M56V-R58Q^). We did so because Chlamydomonas showed its highest sensitivity to rapamycin when expressing the yeast-like CrFKBP12^M52V-E54Q^ mutant (Crespo et al. 2005).

**Fig. 2.**
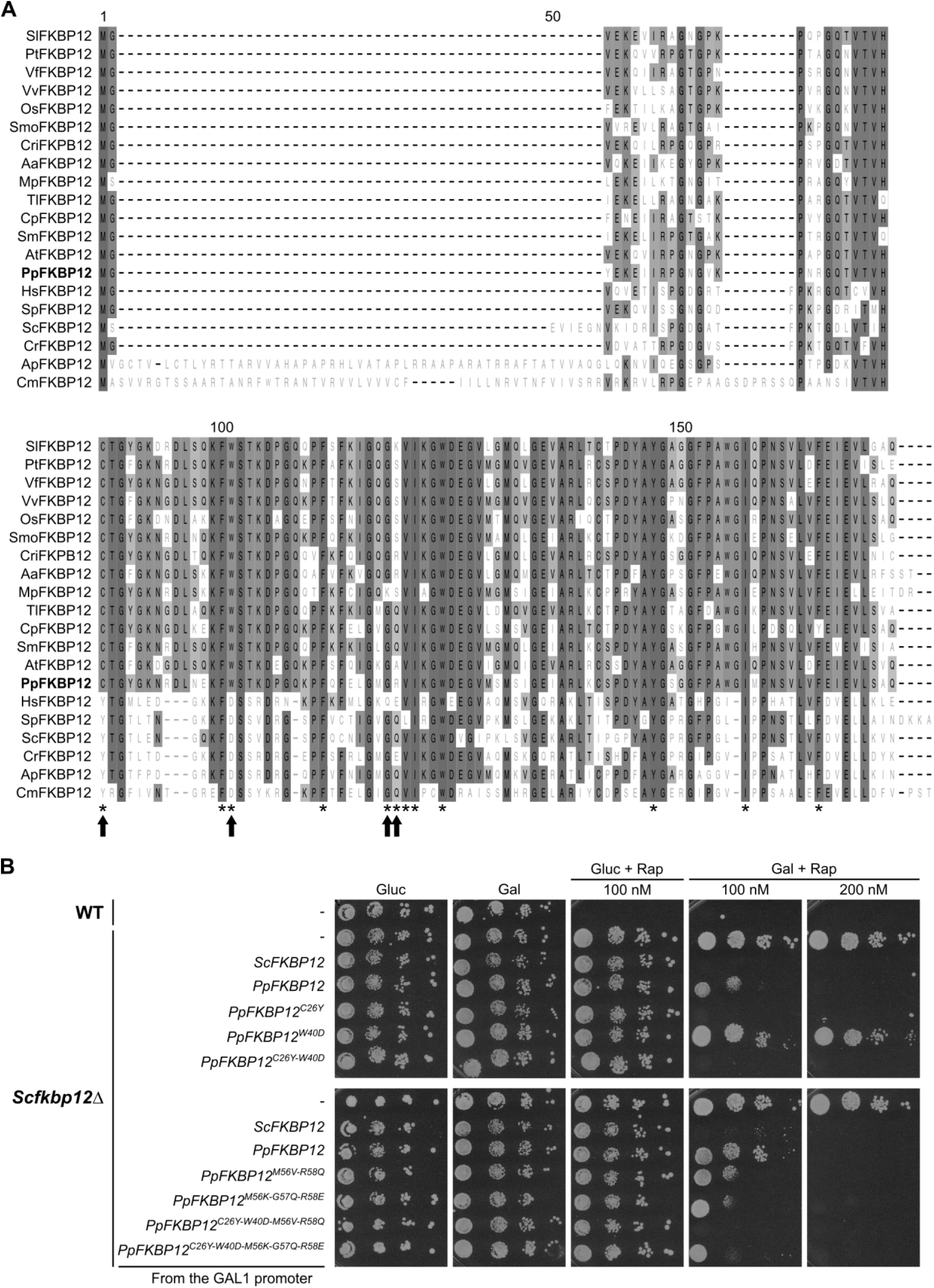
FKBP12 is conserved in Physcomitrella and is able to mediate rapamycin action in yeast. **A** Multiple sequence alignment of FKBP12 proteins from Physcomitrella (bold) and representative organisms: *Anthoceros angustus* (Aa), *Arabidopsis thaliana* (At), *Auxenochlorella pyrenoidosa* (Ap), *Calohypnum plumiforme* (Cp), *Ceratopteris richardii* (Cri), *Chlamydomonas reinhardtii* (Cr), *Cyanidioschyzon merolae* (Cm), *Homo sapiens* (Hs), *Marchantia polymorpha* (Mp), *Oryza sativa* (Os), *Physcomitrium patens* (Pp), *Populus trichocarpa* (Pt), *Saccharomyces cerevisiae* (Sc), *Schizosaccharomyces pombe* (Sp), *Selaginella moellendorffii* (Smo), *Solanum lycopersicum* (Sl), *Sphagnum magellanicum* (Sm), *Takakia lepidozioides* (Tl), *Vicia faba* (Vf), *Vitis vinifera* (Vv). Majority consensus amino acids are shaded in grey. Asterisks indicate residues from human FKBP12 that interact with rapamycin. Arrows indicate amino acids within the rapamycin-binding pocket of HsFKBP12 that are not conserved in PpFKBP12. **B** Yeast complementation assay of the *fkbp12Δ* mutant. Wild-type MH272-3c yeast strain transformed with an empty vector (p415 GAL1), and mutant MH274-1a (*fkbp12Δ*) strain transformed with either p415 GAL1 or p415 GAL1-derived vectors expressing different *FKBP12* variants, were grown in SD-Leu or SGal-Leu suspension cultures. Cultures were normalized to an optical density (OD) of 0.1 at 660 nm, subjected to 10-fold serial dilutions, and 15 µL of each dilution were spotted onto SD-Leu or SGal-Leu plates supplemented with DMSO (0.02%) or with rapamycin at 100 or 200 nM. Plates were incubated at 30°C for 5 days.

To investigate if these mutated PpFKBP12 isoforms exhibit higher affinity to rapamycin than the WT protein, we evaluated their ability to replace ScFKBP12 in a yeast complementation assay. Yeast *fkbp12Δ* cells grow normally in the presence of rapamycin on minimal solid medium containing galactose as a carbon source, unless *ScFKBP12* is ectopically expressed from a plasmid under the galactose-inducible GAL1 promoter (Heitman et al. 1991b; Crespo et al. 2005). We expressed WT *PpFKBP12* or mutant variants driven by the GAL1 promoter in the yeast *fkbp12Δ* strain and spotted them onto galactose plates supplemented with rapamycin. Surprisingly, we found that PpFKBP12 is moderately capable of restoring the sensitivity to rapamycin of yeast *fkbp12Δ* cells (**Figure 2B**). When double the amount of rapamycin was used (200 nM), which is still a relatively low concentration, those cells did not grow at all, which indicated that PpFKBP12 can substantially form a complex with rapamycin and mediate its inhibitory action on TOR, at least in yeast. As for the mutant variants, all but PpFKBP12^W40D^ complemented the rapamycin-resistant phenotype of *fkbp12Δ* cells, and, when growth was evaluated from plates with 100 nM rapamycin, we found that these mutants were more efficient than WT PpFKBP12 in mediating rapamycin action. Intriguingly, yeast expressing PpFKBP12^C26Y^ behaved like those expressing ScFKBP12 (**Figure 2B**). This indicates that having a tyrosine at position 26 instead of a cysteine is required for PpFKBP12 to strongly bind rapamycin. The importance of this tyrosine is highlighted by its ability to reverse the effect of the W40D mutation, and to further enhance the affinity to rapamycin of PpFKBP12^M56K-G57Q-R58E^ and PpFKBP12^M56V-R58Q^, even in the presence of the W40D mutation (**Figure 2B**). Asp38 in HsFKBP12 is involved in a hydrogen bond with rapamycin (Choi et al. 1996), and therefore, it was unexpected that the W40D substitution in PpFKBP12 results in a loss-of-function mutant. The effect of the other substitutions correlates with their role in establishing the bond with rapamycin in human FKBP12.

Taken together, FKBP12 is conserved in Physcomitrella and is able to mediate rapamycin action in yeast *fkbp12Δ* cells. However, despite the fact that certain aa substitutions in PpFKBP12 enhanced its affinity to the drug, these results could not fully explain why Physcomitrella is highly insensitive to rapamycin. Since PpFKBP12 could significantly inhibit yeast growth in the presence of rapamycin, it is very likely that a low expression level of endogenous *FKBP12* is a reason for rapamycin insensitivity in Physcomitrella.

### A rapamycin-sensitive Physcomitrella mutant

A Physcomitrella mutant with acquired sensitivity to rapamycin will enable the functional characterization of PpTOR. Therefore, we engineered transgenic lines that ectopically express *PpFKBP12-cMyc* or *PpFKBP12^C26Y-^cMyc* under the control of the CaMV 35S promoter and terminator from a construct that contains DNA sequences homologous to the PIG1 locus, which enables targeted and stable integration into the *P. patens* genome (Okano et al. 2009). Since Arabidopsis plants overexpressing *HsFKBP12* or *ScFKBP1*2 showed more sensitivity to rapamycin than those overexpressing the endogenous gene (Ren et al. 2012; Xiong and Sheen 2012), we decided to additionally generate Physcomitrella mutants overexpressing the yeast *ScFKBP12-cMyc* or the human *HsFKBP12-cMyc*.

After transformation of protoplasts, regeneration, and selection on hygromycin, between 102 and 115 different plants per DNA construct were obtained. From these, between 32 and 62 different plants per DNA construct were analysed by PCR for correct integration of the expression cassettes at the PIG1 locus. Between 22% and 31% of these plants showed targeted integration in the *P. patens* genome. In addition to targeted integration, multiple and random integrations can occur (Kamisugi et al. 2006; Hoernstein et al. 2023). Therefore, we employed qPCRs on genomic DNA to determine the copy numbers of the integrated expression cassettes. We aimed to select lines with only a single copy of the integrated construct, or with similar copy numbers. Our screen for lines with one copy of the expression cassettes led to the identification of one transgenic plant (ESPB001) containing *PpFKBP12-cMyc* (**Figure 3A**). Interestingly, the addition of 10 µM rapamycin for 7 days to protonema from this plant in a suspension culture slightly impacted their growth (**Figure 3A**). This indicates that one extra copy of *PpFKBP12* is capable of enhancing the sensitivity of Physcomitrella to rapamycin, and that endogenous levels of PpFKBP12 are not high enough to mediate this response. We expected, however, a higher sensitivity to the drug since *PpFKBP12-cMyc* was expressed from the 35S promoter, which usually provides moderate to strong expression levels in Physcomitrella (Horstmann et al. 2004; Niederau et al. 2024). However, it appears that the 35S-driven expression of *PpFKBP12-cMyc* in Physcomitrella is not strong enough to enable high sensitivity to rapamycin, while the 35S-driven expression of *ScFKBP12* in Arabidopsis conferred insensitivity to the inhibitor (Ren et al. 2012). This hypothesis was validated when we tested the sensitivity to 10 µM rapamycin of transgenic plants with increasing copy numbers of the *PpFKBP12-cMyc* cassette, which in general correlated with an increasing amount of the synthesized protein (**Figure 3**), consistent with previous observations in transgenic Physcomitrella lines (Ruiz-Molina et al. 2022). We observed that the more copies were introduced, the more Physcomitrella became sensitive to rapamycin, with line ESPB003 (>500 copies) showing a drastic decrease in biomass accumulation when compared to the control (**Figure 3**). Starting with a protonema suspension at around 100 mg DW/L, ESPB003 reached, after 7 days, 302 ± 2 mg DW/L in the presence of DMSO versus 148 ± 4 mg DW/L in the presence of rapamycin. On the other hand, plants expressing *PpFKBP12^C26Y-^cMyc* did not gain more sensitivity to rapamycin than those with *PpFKBP12-cMyc* (**Figure 3**; line ESPB002 vs line ESPB006 with equivalent copies of the integration cassettes, 109 ± 19 and 111 ± 8, respectively). This result is in line with our finding that both variants can replace ScFKBP12 in the yeast complementation assay (**Figure 2C**; panel 200 mM rapamycin). As for the lines expressing *ScFKBP12-cMyc* or *HsFKBP12-cMyc*, we saw that rapamycin could only slightly affect their growth (**Figure 3A**). This may be due to an inefficient synthesis of the respective proteins, regardless of how many expression cassettes are present in these lines, because these constructs were not codon-optimized for expression in Physcomitrella (**Figure 3B**).

**Fig. 3.**
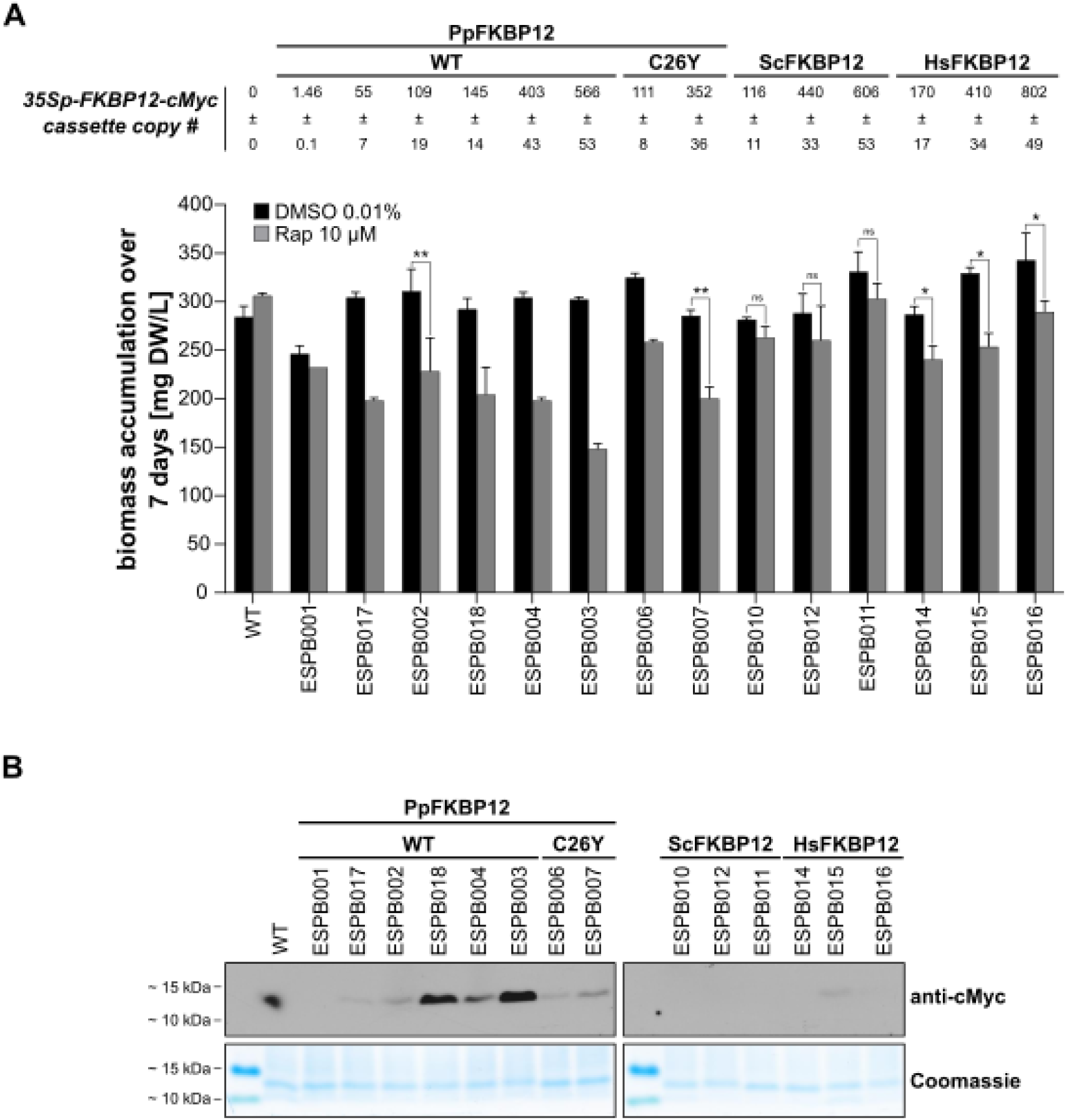
Establishment of a rapamycin-sensitive Physcomitrella line overexpressing endogenous FKBP12. **A** Top. Copy number count of the different *35Sp-FKBP12-cMyc* expression cassettes determined via qPCR in the selected transgenic lines. Bottom. Determination of protonema biomass accumulation over 7 days in the presence of 10 µM rapamycin. Protonema from weekly sub-cultured suspensions of the indicated lines were prepared at a density of 100 mg DW/L in Knop ME pH 5.8. Aliquots of 25 ml were established in 100 mL Erlenmeyer flaks and subjected to treatment with 0.01% DMSO or 10 µM rapamycin. Suspensions were incubated for 7 days under 70 µmol photons m^-2^ s^-1^ at 22 °C in a 16 h/8 h light/dark cycle. Plant material was collected and biomass was determined. Bars represent means of two to three biological replicates; error bars represent standard deviations (SD). Significance levels (*p < 0.05 and **p < 0.01) are based on a paired t-test. NS means not significant. Absence of significance means that statistical power was limited by insufficient replicate number. **B** Protonema of the indicated lines were grown as suspensions in Knop ME pH 5.8. Tissues were harvested, crude protein extracts were prepared, and FKBP12-cMyc was immunoblotted with anti-cMyc antibody. The dot in the WT sample is dirt from the developer and not a band. Protein loading control by Coomassie staining is depicted below.

In conclusion, we generated a Physcomitrella mutant with high levels of PpFKBP12 that was highly sensitive to rapamycin. This suggests that Physcomitrella’s insensitivity to this inhibitor is mainly due to a low expression level of endogenous *PpFKBP12* (**Supplementary Figure S1**). FKBP12 is expressed highest in spores with FPKM (fragments per kilobase of transcript per million mapped reads) values of less than 80 and less than 40 in protonema. These are rather low expression values compared to photosynthesis-related genes, which can have FPKM levels of more than 700 or even beyond 5000 (Niederau et al. 2024). However, rapamycin did not fully arrest the growth of the mutant. This might be due to PpFKBP12 protein levels still not being high enough to mediate a full action of rapamycin. Another plausible explanation is that TORC1 function(s) essential for cell growth and proliferation cannot be fully inhibited by the FKB12-rapamycin-TOR complex, as suggested for the effect of rapamycin on fission yeast (*Schizosaccharomyces pombe*), Arabidopsis, and Chlamydomonas (Weisman et al. 1997; Menand et al. 2002; Crespo et al. 2005). Here, rapamycin drastically inhibited protonema biomass accumulation (**Figure 3A**), an effect also seen in transgenic Arabidopsis seedlings expressing *ScFKBP12* from the 35S promoter with one or several copies of the recombinant expression cassette (Ren et al. 2012). Thus, the observed inhibition may represent the maximal achievable effect of rapamycin on Physcomitrella under these conditions. In conclusion, we found that rapamycin can inhibit Physcomitrella growth, indicating the presence of a functional TOR signalling pathway in this plant.

### Multiple effects of TOR inhibition

To determine whether the observed effect of rapamycin is specific and mediated through PpTOR, rather than resulting from nonspecific toxicity, we evaluated the growth of the strongest *PpFKBP12* overexpressing line in the presence of decreasing concentrations of rapamycin (10 µM, 5 µM, 1 µM). We found that the lower the concentration of rapamycin, the smaller its effect on protonema biomass accumulation is (**Figure 4A**). This supports the conclusion that the effect of rapamycin is specific and likely mediated through PpTOR. Notably, the strongest effect was observed at a concentration of 10 µM. Therefore, this concentration was subsequently used.

**Fig. 4.**
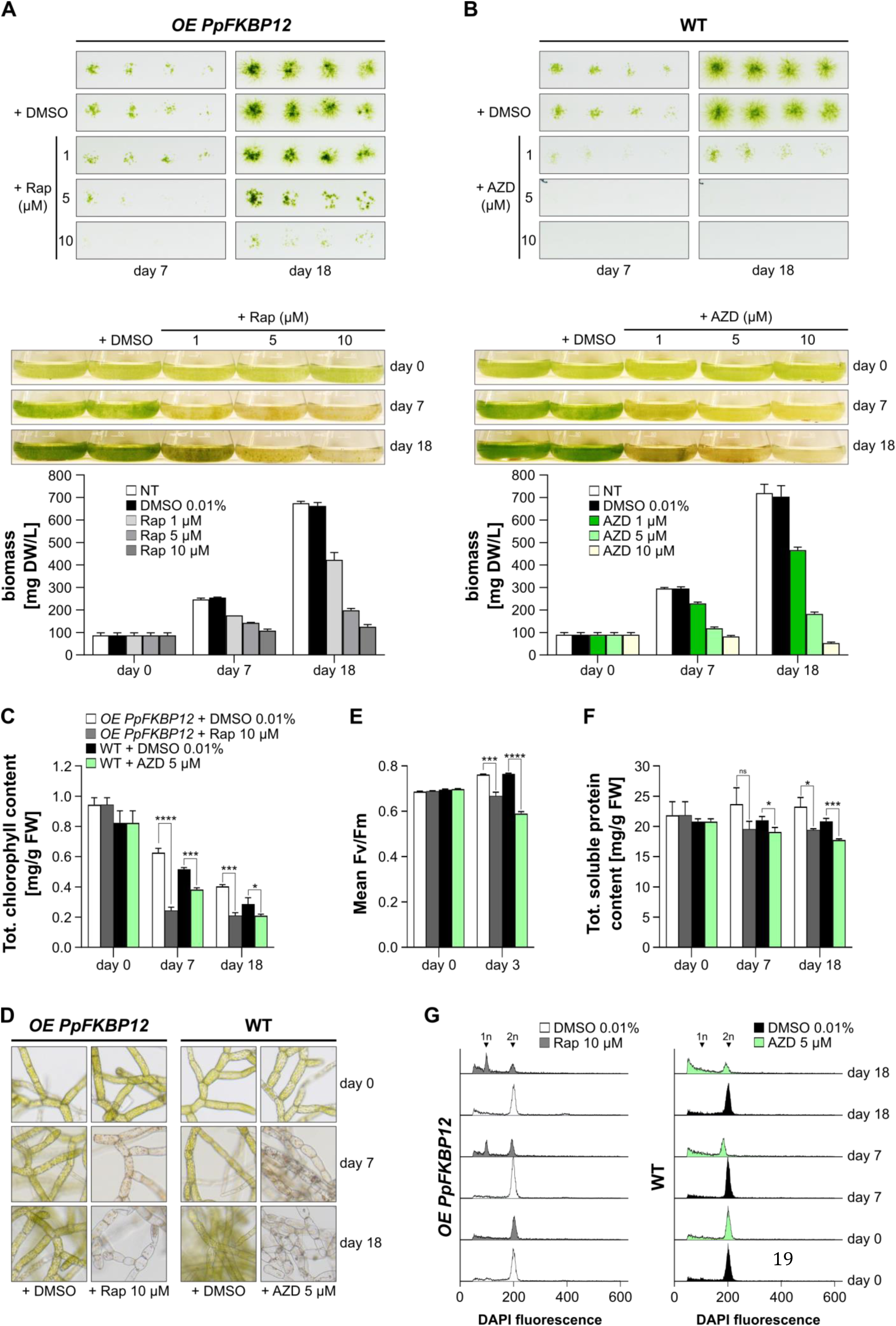
TOR inhibition impacts Physcomitrella growth, induces chlorosis, inhibits photosynthesis, alters total protein content, and delays cell cycle progression. **A** Same experiment as in Figure 1, except that the ESPB003 line overexpressing *PpFKBP12* was analysed. **B** Same experiment as in Figure 1, except that AZD8055 (AZD) instead of rapamycin was used. **C** Chlorophyll content determination in protonema under same growth conditions as in panels A and B at the indicated treatment concentrations. **D** Light microscopy images of protonema under same growth conditions as in panels A and B at the indicated treatment concentrations. **E** Photosynthetic activity measured in protonema under same growth conditions as in panels A and B at the indicated treatment concentrations. Same legend as in panel C. **F** Total protein content in protonema under same growth conditions as in panels A and B at the indicated treatment concentrations. Same legend as in panel C. **G** Histograms of flow cytometric analyses of DAPI stained isolated nuclei of protonema under same growth conditions as in panels A and B at the indicated treatment concentrations. The abscissa represents the channel numbers corresponding to the relative fluorescence intensities of analysed particles (liner mode), while the ordinate indicates the number of counted events (∼5000 counts). 1n and 2n refer to DNA content. Significance levels (*p < 0.05, **p < 0.01, and ***p < 0.001, and ****p < 0.0001) are based on multiple unpaired t-tests, each performed on data from three independent biological replicates. NS means not significant.

Apart from rapamycin, mTOR and AtTOR can be inhibited by a set of chemicals that act as ATP-competitive inhibitors, such as AZD8055 (Chresta et al. 2010; Montané and Menand 2013). In Arabidopsis, AZD8055 is highly specific to the TOR kinase (Perdoux et al. 2024). Unlike rapamycin, AZD8055 achieves a more complete inhibition of TOR (Van Leene et al. 2019; Liu et al. 2024), which could be useful to elucidate TOR functions that are not affected by a treatment with rapamycin. Therefore, we additionally evaluated the effects of AZD8055 on Physcomitrella protonema biomass accumulation. Similar to rapamycin, we recorded a dose-dependent response, with 10 µM of AZD8055 completely blocking protonema growth (**Figure 4B**). These results suggest that AZD8055 can also be applied to study TOR action in Physcomitrella, and may be best suited if a complete inhibition of the signalling pathway is required.

Whether with rapamycin or AZD8055, protonemata turned yellow under TOR inhibition (**Figure 4**), an effect seen in other photosynthetic organisms as well (Caldana et al. 2013; Upadhyaya and Rao 2019). We measured total chlorophyll contents in the Physcomitrella line overexpressing *PpFKBP12* before and after the addition of 10 µM rapamycin, and found a significant decline when compared to the control (**Figure 4C**). The same was observed when 5 µM AZD8055 was applied to WT protonema (**Figure 4C**). TOR is crucial for regulating chlorophyll synthesis, and chloroplast development and function (Couso et al. 2021; Liu and Xiong 2022; D’Alessandro et al. 2024). Light microscopy revealed that chloroplasts had shrunk to such an extent that they were difficult to spot in some cells (**Figure 4D**), indicating that TOR inhibition affects shape and abundance of chloroplasts in Physcomitrella, which can explain the observed decrease in chlorophyll levels. This decline likely includes autophagy, as TOR is a central regulator of autophagy in plant energy stress responses, and its inhibition ultimately leads to cell death (Neufeld 2010; Feng et al. 2025).

Unexpectedly, we also observed a progressive decline in chlorophyll content between day 0 and days 7/18 in the controls which were treated with DMSO (**Figure 4C**). This decline cannot be attributed to changes in chloroplast morphology or quantity in chloronema cells, the juvenile protonemal cells predominant at day 0. Instead, the differentiation of chloronema into caulonema, which are elongated cells with fewer chloroplasts (Reski and Cove 2004), likely explains the reduced chlorophyll levels. Consistent with this, caulonema formation increased remarkably by days 7 and 18 as compared to day 0 (**Supplementary Figure S2**).

In Chlamydomonas and Arabidopsis, TOR inhibition impacts chloroplast functions (Couso et al. 2021; D’Alessandro et al. 2024). This can be assessed by measuring the maximal efficiency of photosystem II (PSII), which reflects the photosynthetic activity of a plant and is not affected by variations in chlorophyll levels. To test whether chloroplast functions in Physcomitrella are affected by TOR inhibition, we measured chlorophyll fluorescence in protonema treated with rapamycin or AZD8055 and calculated the PSII maximum quantum yield (Fv/Fm). Treatment of protonema with rapamycin or AZD8055 resulted in a drop of the photosynthetic efficiency. In protonema of *OE PpFKBP12*, we measured an Fv/Fm of 0.763 ± 0.002 with 0.01% DMSO vs 0.667 ± 0.017 with 10 µM rapamycin after 3 days of treatment. With 5 µM AZD8055, WT protonema showed an Fv/Fm value of 0.591 ± 0.008 vs 0.766 ± 0.002 for those treated with 0.01% DMSO (**Figure 4E**).

Next, we investigated if Physcomitrella TOR also governs other cellular mechanisms such as protein homeostasis and cell division, as reported for other organisms (Liu and Xiong et al. 2022; Liu et al. 2025). We found that TOR inhibition led to a drop in cellular total soluble protein content (**Figure 4F**) and to delays in cell cycle progression (**Figure 4G**). The addition of rapamycin resulted in chloronema cells accumulating in the G1 phase of the cell cycle, similar to observations in yeast (Barbet et al. 1996), whereas under normal conditions, Physcomitrella chloronema is congregated at the G2/M transition (Schween et al. 2003). AZD8055 treatment of protonema did not result in the formation of a pronounced peak of cells arrested at G1. However, it substantially reduced the number of cells accumulating with duplicated DNA content and increased the counted events with a fluorescence intensity below 200, thus suggesting cell cycle arrest (**Figure 4G**).

Collectively, we found that TOR signalling is crucial for regulating chlorophyll synthesis and chloroplast development and function, protein synthesis, and cell cycle progression in Physcomitrella, underscoring its evolutionary conserved function in plants.

### Physcomitrella encodes TOR, LST8 and RAPTOR

In yeast and mammals, the TOR kinase operates in distinct cellular complexes with different functions, namely TORC1 and TORC2. Therefore, our next step was to determine whether the main protein components of these complexes exist in Physcomitrella and whether they have maintained conserved functions. Using the Arabidopsis peptide sequences of TOR, LST8-1, and RAPTOR1 as queries, a BLASTP analysis revealed the presence of single genes in the Physcomitrella genome that code for putative homologs of AtTOR and AtLST8-1, whereas four different putative PpRAPTOR proteins are encoded. These genes are *Pp3c6_4910* and *Pp3c11_25160* for PpTOR and PpLST8, respectively. Those encoding the PpRAPTOR proteins are *Pp3c9_4770*, *Pp3c25_7220*, *Pp3c17_3950* and *Pp3c15_3360* (hereafter referred to *PpRAPTOR1*, *PpRAPTOR2, PpRAPTOR3, and PpRAPTOR4*, respectively). Phylogenetic reconstruction revealed frequent, but lineage-specific duplications of *RAPTOR* genes throughout land plants (**Supplementary Figure S3**). The expansion in Physcomitrella is fully shared by one-to-one orthologs in *Funaria hygrometrica* – a moss from the same family – is partially shared by three orthologs in *Ceratodon purpureus* – a moss from a different subclass – and is not shared by *Takakia lepidozioides*, which is sister to all other mosses. Interestingly, the *RAPTOR* genes of Physcomitrella are located on four different chromosomes. And while *C. purpureus* encodes direct orthologs of both PpRAPTOR2 and PpRAPTOR3, Physcomitrella and *F. hygrometrica* share an additional duplication that occurred after the split of their common ancestor from *C. purpureus*. Since Physcomitrella and *F. hygrometrica* share two rounds of whole genome duplications (WGDs), with the older event being shared with *C. purpureus* (Carey et al. 2021) and the younger event only predating the origin of the Funariaceae (Lang et al. 2018; Kirbis et al. 2025), the four genes of Physcomitrella might be remnants of both WGDs. A structural domain analysis reveals the presence of all known functional domains within these proteins (**Figure 5A**). Publicly available transcription data indicates that they are expressed at low to moderate levels similar to FKBP12 across all tissue types (**Supplementary Figure S1**).

**Fig. 5.**
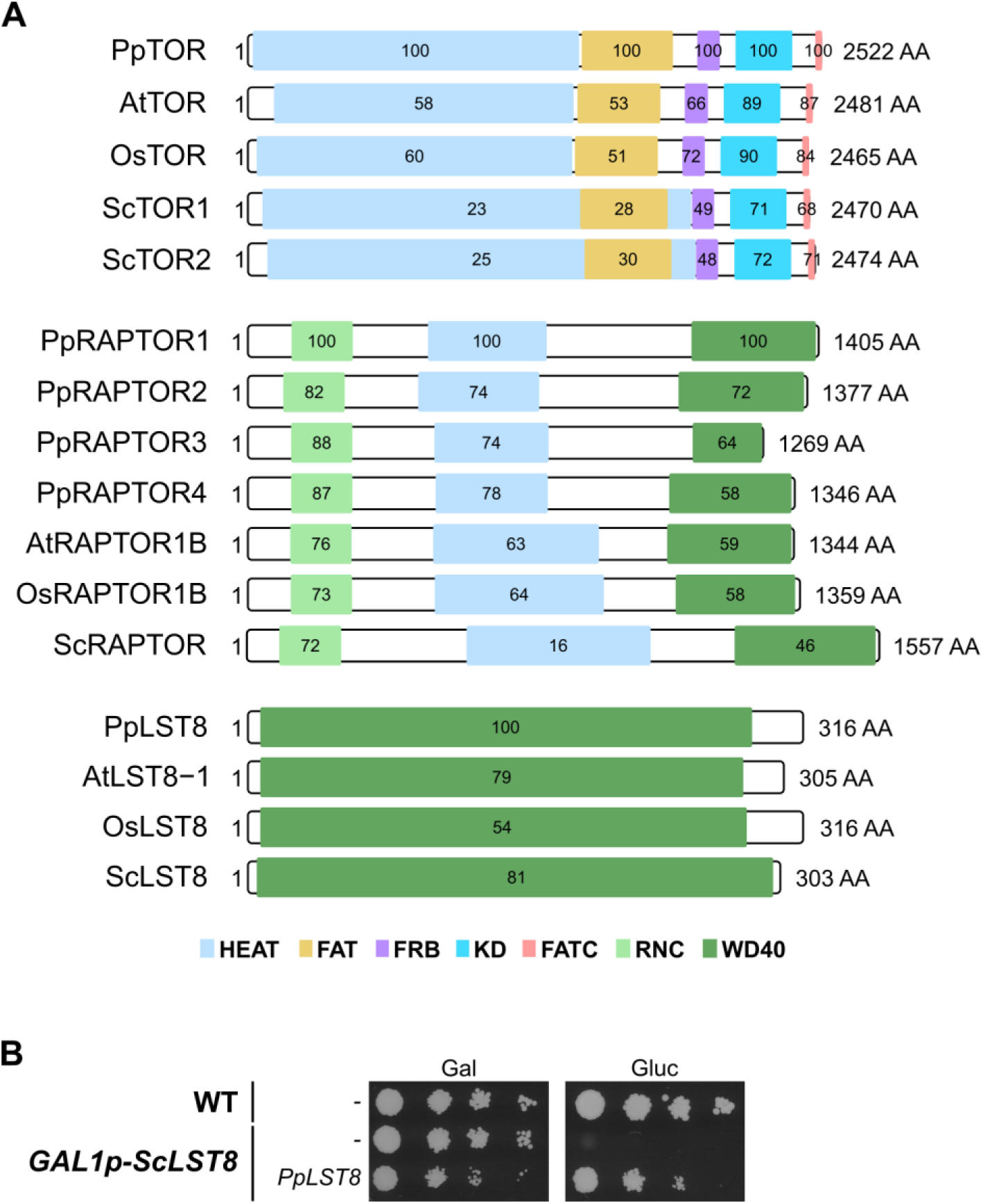
TOR, LST8 and RAPTOR are conserved in Physcomitrella. **A** Conserved protein domains of PpTOR, PpLST8, or the four PpRAPTOR paralogs with homologs from *A. thaliana* (At), *O. sativa* (Os), *S. cerevisiae* (Sc). Values in domain boxes indicate percentage of shared identity with the respective homologous Physcomitrella domain. **B** Complementation of a yeast *lst8* mutant with the Physcomitrella *LST8* cDNA. Wild-type TB50a yeast strain and mutant RL57-2d expressing *ScLST8* from the galactose-inducible GAL1 promoter were initially grown as suspensions in rich medium containing galactose as a carbon source (YPGal). Strains were transformed with pESPB019 (empty vector) or pESPB020 (pESPB019-derived vector expressing *PpLST8*) and selected on SGal-Leu plates. Selected transformants were grown for 6 h at 30°C in SD-Leu suspension cultures. Cultures were normalized to an OD (660 nm) of 0.2, subjected to 10-fold serial dilutions, and 15 µL of each dilution were spotted onto SGal-Leu or SD-Leu plates, which were incubated at 30°C for 5 days. Plates were scanned upside down with the lid open on a Canon CanoScan 9000F scanner.

Whereas Arabidopsis and Physcomitrella encode one *TOR* gene each, Physcomitrella encodes one *LST8* gene and Arabidopsis two. The difference is more pronounced with the *RAPTOR* genes, of which Arabidopsis encodes two and Physcomitrella four. This expansion indicates neofunctionalisation of the *RAPTOR* genes in Physcomitrella.

For a functional analysis, we tested whether the Physcomitrella genes could substitute their yeast homologs in promoting yeast cell growth. In yeast, *TOR*, *LST8* and *RAPTOR* are essential genes (Loewith et al. 2002). Therefore, we used three well-described yeast strains in which the expression of the respective genes could be shut down by shifting the cells from galactose to glucose. Those strains are SW137-4b, RL57-2d, and RL93a (Loewith et al. 2002; Wullschleger et al. 2005; Díaz-Troya et al. 2008). *S. cerevisiae* has two *TOR* genes, *TOR1* and *TOR2*. TORC1 contains either one of the proteins, but TORC2 contains exclusively TOR2. However, the formation of both complexes is essential for the viability of yeast cells. Therefore, SW137-4b lacks *TOR1* and expresses *TOR2* from the galactose-inducible GAL1 promoter. In RL57-2d and RL93a, *LST8* and *RAPTOR* were placed under the GAL1 promoter, respectively. We introduced plasmids expressing the respective Physcomitrella full-length cDNA under the strong and constitutive GPD (Glyceraldehyde-3-Phosphate Dehydrogenase) promoter into these yeast strains and analysed their growth on synthetic minimal plates, containing either galactose or glucose as the carbon source. A WT TB50a yeast strain and mutants containing empty vectors served as positive and negative controls, respectively, for growth on plates with glucose.

Surprisingly, only PpLST8 could complement its respective yeast mutant (**Figure 5B**, **Supplementary Figure S4**), revealing that PpLST8 is a functional homolog of ScLST8. This also indicates that PpLST8 can bind ScTOR and ScRAPTOR to form a complex in yeast. In contrast, none of the four Physcomitrella RAPTORs could substitute for its yeast homolog. This is not due to the use of an incorrect strain, since the introduction of a plasmid expressing ScRAPTOR restored yeast growth in the presence of glucose (**Supplementary Figure S4**). Likewise, PpTOR could not complement its yeast homolog. Here, the low degree of conservation of the HEAT and FAT domains with 23% and 28% sequence identity, respectively (**Figure 5A**), could result in improper assembly, function, and localization of ScTORC. This hypothesis is supported by the findings that rice TOR (OsTOR) could not rescue yeast *tor* mutants either, but a chimeric TOR protein with the yeast HEAT and FAT domains and the rest from OsTOR could (Maegawa et al. 2025). In yeast, RAPTOR interacts with its C-terminal WD40 domains with the HEAT repeats of TOR, whereas its N-terminus is positioned next to the TOR kinase domain, likely bringing substrates into the vicinity of the catalytic region (Adami et al. 2007). As the WD40 domains of ScRAPTOR and PpRAPTOR1 share 46% of sequence identity, and the HEAT repeats share only 16% of sequence identity (**Figure 5A**), it is likely that the domains of PpRAPTOR1 are not conserved enough to allow biogenesis of ScTORC1. So far, only OsRAPTOR2 has been able to support growth of a thermosensitive yeast *raptor* mutant (Maegawa et al. 2025). This assay was conducted at 37°C, a temperature that could strengthen the interaction between OsRAPTOR2 and ScTOR, which under normal growth conditions is very weak, if not absent. We did not find homologs of key TORC2 proteins, such as RICTOR and mSIN1, in Physcomitrella, which is in line with the findings in other photosynthetic organisms.

In summary, Physcomitrella LST8 is functionally conserved and complements its yeast homolog, whereas neither Physcomitrella TOR nor its four RAPTOR proteins rescued the respective yeast mutants. Despite conserved domain architectures, critical sequences or structural divergences, particularly in regions mediating protein-protein interactions, may prevent proper assembly and function of TORC1 in yeast. Further experiments like the one conducted with yeast/rice chimeric TOR could confirm this.

### Downregulation of TOR, LST8 or RAPTOR delays development

Next, we investigated whether a reduced expression of PpTORC1 subunit genes impacts not only Physcomitrella growth, but also its development. Specifically, we evaluated whether TOR activity influences the developmental transition from two-dimensional (2D) to three-dimensional (3D) growth in Physcomitrella, also known as budding (Reski and Abel 1985). Budding is the first key developmental decision in the life cycle of mosses (Reski 1998). Moreover, this 2D to 3D transition is a key developmental innovation of land plants, and thus distinguishes algae from plants (Weeks et al. 2025). To investigate this, WT protonemal tissue derived from suspension culture was transferred onto solid Knop ME media supplemented with either 0.02% glucose or 1 µM AZD8055 as described in Hoernstein et al. (2023). Plates without any supplementation served as controls. The formation of leafy gametophores was subsequently monitored over time. In the presence of 1 µM AZD8055, gametophore development was markedly suppressed throughout the observation period. In contrast, the number of gametophores began to increase noticeably from day 12 onwards in the control condition, with an even more pronounced promotion observed on plates containing 0.02% glucose (**Figure 6A**). The same effect was observed when employing 1% sucrose instead of 0.02% glucose (**Supplementary Figure S5**). This demonstrates that inhibition of TOR activity in Physcomitrella not only impairs growth (**Figure 4**) but also development (**Figure 6A**). Conversely, supplementation with glucose or sucrose, which are known to promote TOR activity in plants and mammals, promotes developmental progression in Physcomitrella. Likewise, sugar availability affects developmental progression in Physcomitrella via cyclin D (Lorenz et al. 2003), suggesting a feedback loop between nutrients, TOR activity, cyclin D expression, and the cell cycle in Physcomitrella.

**Fig. 6.**
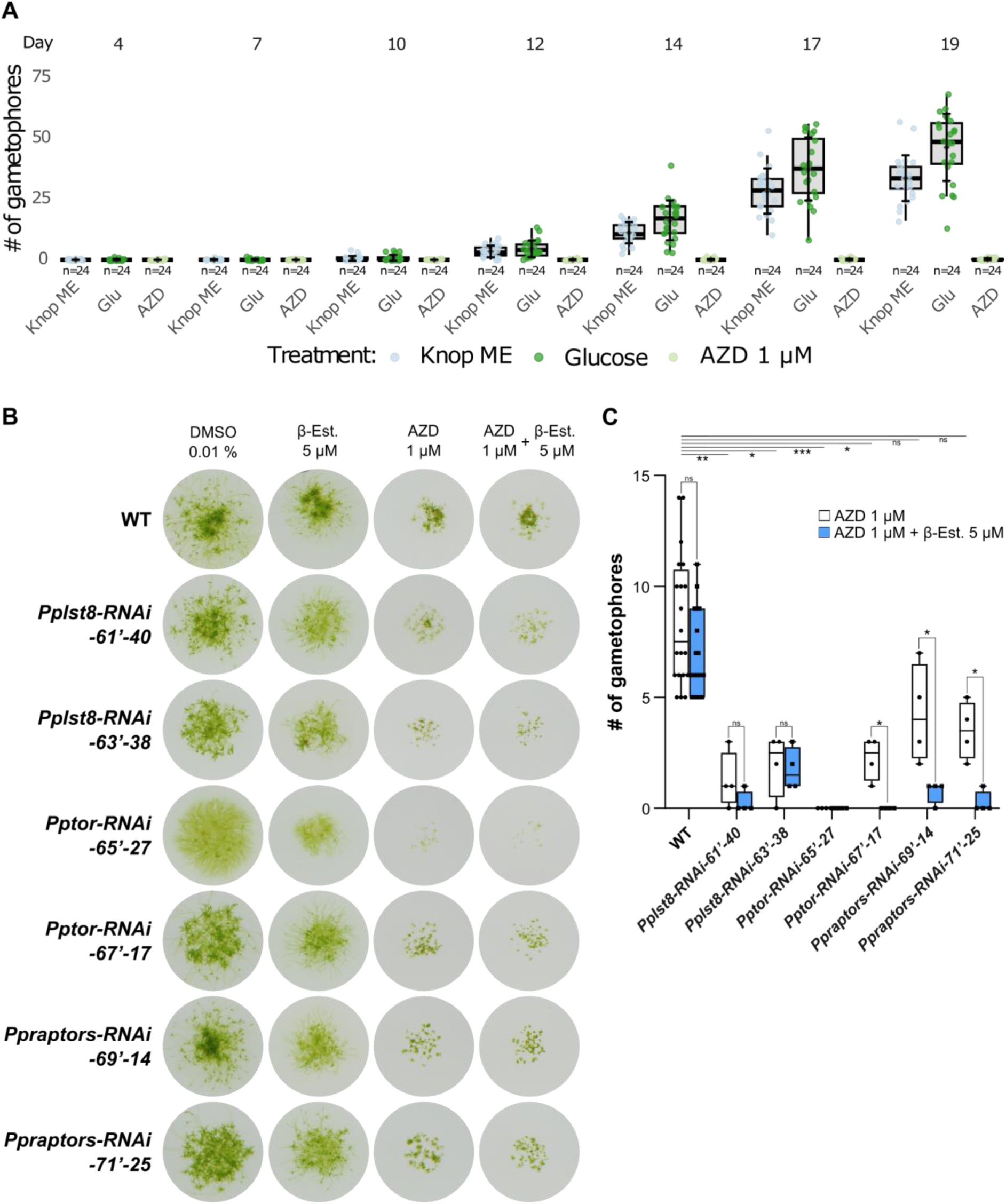
Downregulation of TOR, LST8 or RAPTOR delays Physcomitrella development. **A** Box plot showing the number of gametophores per colony, counted after transferring 25-day-old WT protonema on Knop ME agar plates, or plates supplemented with 0.02% glucose or 1 µM AZD8055 (n = 24). Data are represented as mean +-SD. 15 µL droplets at a density of 440 mg DW/L were spotted. **B** Weekly sub-cultured protonema in suspension of wild-type, ESPB020 (*Pplst8-RNAi-61’-40*), ESPB024 (*Pplst8-RNAi-63’-38*), ESPB027 (*Pptor-RNAi-65’-27*), ESPB032 (*Pptor-RNAi-67’-17*), ESPB035 (*Ppraptors-RNAi-69’-14*), and ESPB040 (*Ppraptors-RNAi-71’-25*) lines were prepared at a density of 100 mg DW/L in Knop ME pH 5.8. One-fold serial dilutions were conducted and 50 µL were spotted on solid Knop ME pH 5.8 plates supplemented with 0.01% DMSO, 5 µM β-Estradiol (β-Est.), 1 µM AZD8055, or with 5 µM β-Estradiol and 1 µM AZD8055. Plates were photographed after 4 weeks of incubation under 70 µmol photons m^-2^ s^-1^ at 22 °C in a 16 h/8 h light/dark cycle. **C** Box plot showing the number of gametophores per colony of the indicated lines, after 4 weeks of incubation on solid Knop ME pH 5.8 plates supplemented with 1 µM AZD8055, or, with 5 µM β-Estradiol and 1 µM AZD8055. Counting was performed on 19 wild-type colonies, and on 4 colonies of each of the transgenic lines. The box plots depict the mean (horizontal line within the box) of the data, the interquartile range (box), and the 1.5x interquartile range (whiskers). The significance level (*p < 0.05, **p < 0.01, and ***p < 0.001, and ****p < 0.0001) between the AZD8055-treated WT line and each of the AZD8055-treated RNAi lines is based on a one-way ANOVA followed by a post-hoc multiple comparison test against the WT group. A multiple unpaired t-test, with correction for multiple comparisons using the False Discovery Rate (FDR) method set at 0.05, was used to determine statistical significance (*p < 0.05, **p < 0.01, and ***p < 0.001, and ****p < 0.0001) between AZD8055 and AZD8055 + β-Estradiol-treated samples; ns: not significant.

Next, we generated Physcomitrella mutants with estradiol-inducible RNA interference (RNAi), targeting transcripts of individual genes encoding PpTORC1 subunits. This inducible approach was chosen to avoid experimental difficulties associated with the essential nature of these genes, as reported for other organisms (Loewith et al. 2002; Menand et al. 2002; Deprost et al. 2005; Moreau et al. 2012). Additionally, by designing the RNAi construct to target conserved regions, it is possible to attenuate the four PpRAPTOR paralogs simultaneously.

The screen of 60 hygromycin-resistant clones on solid Knop ME plates containing estradiol led to the identification of at least one transgenic line for each of the RNAi constructs, that showed an altered developmental phenotype, reflected by the reduced number and size of the gametophores (**Figure 6B**). One particular line, *Pptor-RNAi-65’-27*, which expresses an RNAi construct targeting the 500-bp coding region (561-1060) of *PpTOR*, equivalent to the one used to generate the Arabidopsis estradiol-inducible *tor* RNAi lines (Xiong and Sheen 2012), did not develop any gametophores on the estradiol-containing plate, whereas only few were spotted on the control plate containing DMSO (**Figure 6B**). This altered differentiation of the *Pptor-RNAi-65’-27* line observed in the absence of estradiol could be due to leakage in the estradiol-inducible RNAi system, as such issues have been reported previously (Nakaoka et al. 2012). Off-target effects could also account for this phenotype. However, we sought to reduce this likelihood by isolating distinct RNAi mutant lines for each gene, where in each line a different region within the transcript sequence is being targeted. We observed consistent phenotypes between these lines (**Figure 6B**), indicating that the developmental phenotypes observed here result from the knock-down of these genes.

It is well documented that organisms with low TOR signalling, including plants, are generally more susceptible to TOR inhibitors, such as rapamycin and AZD8055, at low concentrations (Montané et al. 2013; Jordan et al. 2014; Saliba et al. 2018). This increased sensitivity facilitates screens for mutants with reduced TOR activity. Since the Physcomitrella lines exhibited slightly altered differentiation under control conditions, we grew them on plates containing 1 µM AZD8055, which still allows growth (**Figure 4B**). If the RNAi systems in these lines are leaky, one would expect them to develop fewer gametophores than WT, which was the case (**Figure 6B**). Remarkably, the *Pptor-RNAi-65’-27* line barely grew. The altered differentiation of the other lines was more evident when quantified by counting gametophores. With line *Ppraptors-RNAi-69’-14* being the only exception, the presence of AZD8055 reduced the number of gametophores per colony (WT 8.45 ± 2.83, *Pplst8-RNAi-61’-40* 1.25 ± 1.09, *Pplst8-RNAi-63’-38* 2.00 ± 1.22, *Pptor-RNAi-65’-27* 0.00 ± 0.00, *Pptor-RNAi-67’- 17* 2.25 ± 0.83, *Ppraptors-RNAi-69’-14* 4.25 ± 1.92, and *Ppraptors-RNAi-71’-25* 3.50 ± 1.12, **Figure 6C**), suggesting a reduced TOR signalling in these transgenics, most likely because of RNAi leakage. Addition of estradiol to the AZD8055-containing plates further reduced the number of gametophores in the majority of the analysed colonies, indicating that the estradiol-inducible RNAi systems in these lines were triggered further (**Figure 6C**). The bud numbers did not deviate much between untreated WT, plants on DMSO and beta-estradiol after 4 weeks. Together, the knockdown of *PpTORC1* subunit genes reduces the number of gametophores per colony, highlighting the importance of TOR signalling for developmental progression in Physcomitrella, which is in line with our observations under TOR activity inhibiting conditions.

### TOR-dependent phosphorylation of RPS6 on Ser241

In response to environmental cues, TOR triggers cellular processes through phosphorylation of several downstream effector proteins. The phosphorylation sites within these effectors serve as markers of TOR activity in various species, typically by generating antibodies specific to the phosphorylated form and detecting them in Western blots (Xiong and Sheen 2012; Dobrenel et al. 2016; Saliba et al. 2018). This is an effective tool to investigate TOR signalling at the cellular level which we attempted to implement for Physcomitrella.

However, our attempts to detect phosphorylated ribosomal S6 kinase in Physcomitrella with an antibody against phosphorylated Thr389 in HsS6K1, which also recognizes phosphorylated Thr449 and Thr455 in Arabidopsis S6K1 and S6K2, respectively (Xiong and Sheen 2012), were not successful. Most likely this is due to their low abundance rather than a lack of conservation of the TOR phosphorylation motif FLGF<u>T</u>YVAP in human S6K1 (**Supplementary Figure S6**). This hypothesis is consistent with findings in Arabidopsis, where successful detection with this antibody often requires transgenic plants overexpressing AtS6K1 (Xiong and Sheen 2012; Primo et al. 2022). We therefore shifted to assaying the phosphorylation of RPS6, which is phosphorylated by S6K in a TOR-dependent manner. In Physcomitrella, four *RPS6* genes exist and their expression levels are up to 5-fold higher than those of S6K (**Supplementary Figure S7**). Despite the overall high sequence identity of the Physcomitrella RPS6 isoforms with RPS6A from Arabidopsis (**Figure 7A**), the sequence window around the known phosphorylation site of AtRPS6A (SRL<u>SS</u>AAAKPSVTA; Dobrenel et al. 2016) is far less conserved (**Figure 7B**). Consequently, the commercially available antibody against phosphorylated Ser240 in AtRPS6A could not be used (Agrisera, personal communication).

**Fig. 7.**
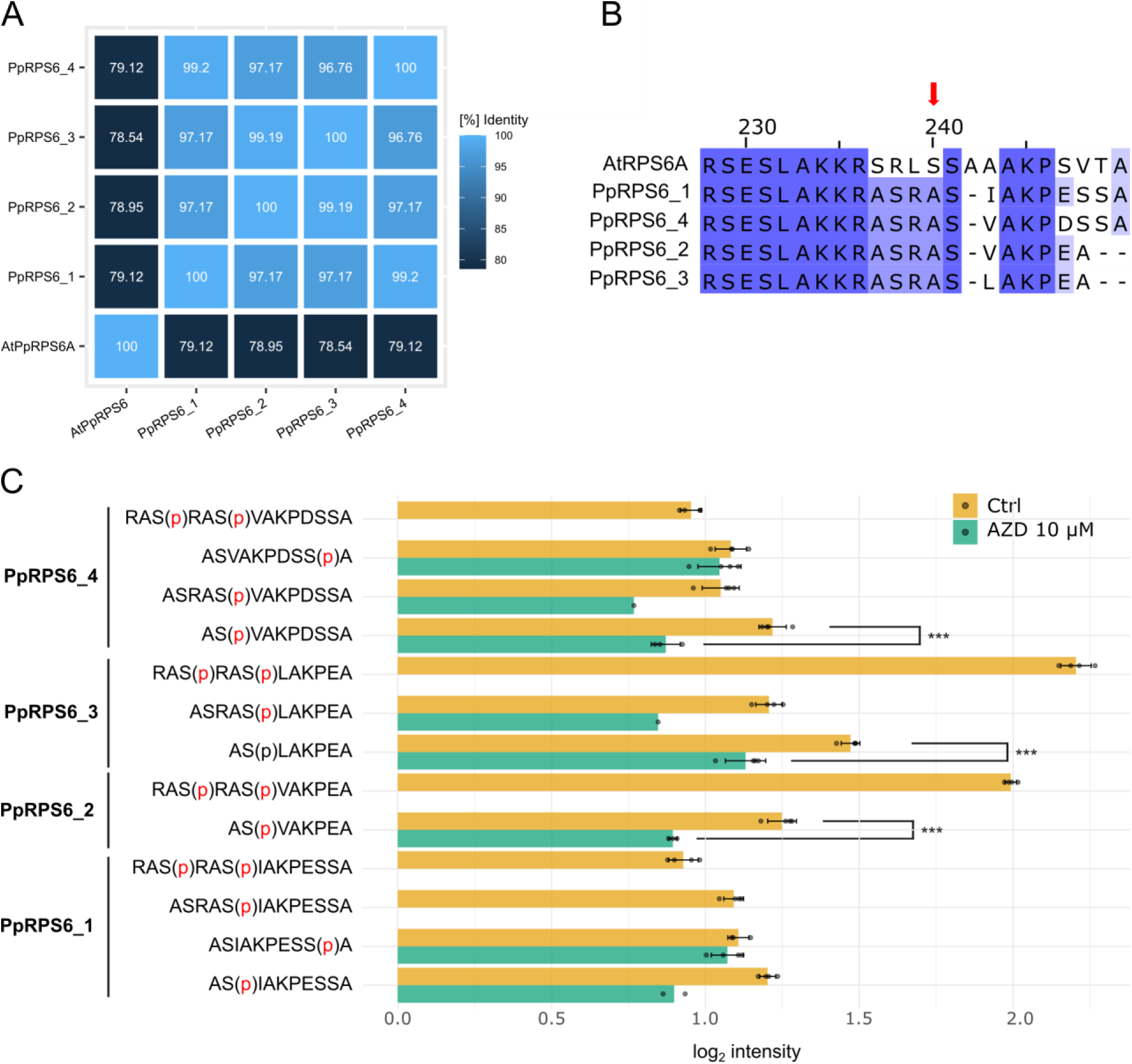
Sequence identity between Physcomitrella and Arabidopsis RPS6 isoforms and analysis of TOR-dependent phosphorylation of Physcomitrella RPS6. **A** Percentage identity matrix of Physcomitrella RPS6 isoforms (PpRPS6_1: Pp3c11_21120V3.1; PpRPS6_2: Pp3c15_25430V3.1; PpRPS6_3: Pp3c15_25780V3.1; PpRPS6_4: Pp3c7_5780V3.1) with Arabidopsis AtRPS6A (AT4G31700.1). Sequences were aligned using *Clustal Omega* (V2.1, Sievers and Higgins 2018) on UNIPROT (https://www.uniprot.org/align). **B** Section of a multiple sequence alignment of Physcomitrella RPS6 isoforms with Arabidopsis AtRPS6A. Sequences were aligned in *Jalview* (2.11.4.1, Waterhouse et al. 2009) using *Mafft* with defaults (Katoh and Standley 2013). Serine 240 in AtRPS6A that is phosphorylated in a TOR-dependent manner is indicated with a red arrow. **C** Phosphorylation level of the Physcomitrella RPS6 C-terminal peptides following the addition of AZD8055 for 60 min. Bars represent mean median normalized intensity values with standard deviation. Each datapoint represents the intensity sum for a peptide per replicate (including different charge states). Significance levels (***p<0.001) are based on a Welch test. The Welch test was only calculated for peptides with at least three data points (biological replicates).

Therefore, we aimed to establish a mass spectrometry-based assay to determine which residues in PpRPS6 undergo phosphorylation in a TOR-dependent manner. Whereas in Arabidopsis two consecutive serines (Ser240 and Ser241) exist, Physcomitrella RPS6 proteins have only a single serine residue (Ser241). We collected protonema from suspensions treated with 10 µM AZD8055 for 60 min, alongside DMSO-treated controls, enriched phospho-peptides by Ti-NTA affinity chromatography and identified the enriched phospho-peptides by LC-MS/MS in data-independent acquisition (DIA) mode. We identified peptides spanning the potential phosphorylation site of interest (around Ser241) for all four PpRPS6 proteins. Among those peptides, Ser241 was identified to be phosphorylated in all four isoforms. Two additional phosphorylation sites were detected, namely Ser238 and Ser248, whereby Ser248 was only found in PpRPS6_1 and PpRPS6_4 (**Figure 7C**). Of those three sites, only phosphorylation of Ser241 showed a significant decrease after 60 min treatment with AZD8055. This effect was observed in the three isoforms PpRPS6_2, PpRPS6_3, and PpRPS6_4. Although a similar trend was noted for PpRPS6_1, the number of data points was insufficient for statistical analysis in this isoform (**Figure 7C**).

Taken together, Physcomitrella TOR mediates PpRPS6 phosphorylation at Ser241. This finding reveals that another key component of the TOR signalling pathway, well characterized in other eukaryotes, is conserved in Physcomitrella.

## Discussion

The signalling hub TOR is studied in algae and flowering plants, yet its role in non-vascular plants remained less clear. Here, we present evidence of TOR signalling in the non-vascular moss plant Physcomitrella. In yeast and animals, TOR signalling is inhibited by rapamycin. Due to the essential nature of the *TOR* gene, rapamycin sensitivity has been key for dissecting TOR function in these organisms. However, assessing this in photosynthetic organisms has been challenging, as most of them are insensitive to rapamycin. This has been attributed to an FKBP12 protein lacking critical amino acids needed to form an effective ternary complex with rapamycin and TOR, a low cellular abundance of FKBP12, or a combination of both. Similar to other plants, we found that Physcomitrella is not sensitive to rapamycin, primarily due to low expression of the *PpFKBP12* gene. By generating a transgenic line overexpressing *PpFKBP12*, we demonstrated that rapamycin treatment delays protonemal growth and differentiation, induces chlorosis, inhibits photosynthesis, reduces total protein content, and delays cell cycle progression. These phenotypic and physiological changes were similarly induced by treatment with the ATP-competitive TOR inhibitor AZD8055. While no TORC2-specific components such as RICTOR or SIN1 are encoded by the *P. patens* genome, we identified and functionally characterised key TORC1 components, including LST8, RAPTOR, and the downstream target RPS6, which undergoes TOR-dependent phosphorylation at Ser241. These findings reveal that the evolutionary conserved TORC1 pathway is present and functional in Physcomitrella. We infer TOR engagement from convergent pharmacological responses and the decrease in RPS6 Ser241 phosphorylation, recognizing that moss-specific inhibitor specificity remains to be formally validated.

Although overexpression of *PpFKBP12* rendered Physcomitrella sensitive to rapamycin, the yeast complementation assay of the *Scfkbp12Δ* mutant indicated that the sequence of PpFKBP12 is not optimal for full interaction with the inhibitor. While the W40D substitution resulted in a loss-of-function mutant, all other aa changes within the rapamycin-binding pocket, aimed at getting a human- or yeast-like PpFKBP12 variant, resulted in proteins that complemented the *Scfkbp12Δ* strain better than WT PpFKBP12. One mutant, PpFKBP12^C26Y^, exhibited an ScFKBP12-like function capacity. The presence of a tyrosine instead of a cysteine at position 26 in PpFKBP12 appeared to be important for efficient rapamycin binding as it reversed the phenotype of the W40D substitution. What could be the reason behind these observations? Unique to land plant FKBP12 proteins are two cysteines at fixed positions (Cys26 and Cys80), reported to be spatially adjacent to potentially form disulphide linkage (Xu et al. 1998), whereas homologs from other organisms contain no or only one cysteine at variable positions. Even though the reduction of this disulphide bridge was shown to decrease the affinity of *Vicia faba* FKBP12 towards its ligands *in vitro* (Xu et al. 1998), it is possible that disruption of this bridge could lead to an increased affinity of FKBP12 *in vivo*. Our complementation assay of the *Scfkbp12Δ* mutant with PpFKBP12^C80I^ indicate that this is the case, at least in PpFKBP12. PpFKBP12^C80I^ was as efficient as PpFKBP12^C26Y^ in inhibiting yeast growth in the presence of rapamycin. However, the effects of these two mutations in Arabidopsis FKBP12 could not be replicated, as only AtFKBP12^C26Y^ showed a slightly increased affinity to rapamycin. This contingency could be due to differences in the aa sequences between PpFKBP12 and AtFFKBP12, which might mask the expected phenotype. Therefore, further experiments are needed to validate the above hypothesis. For instance, by combining the C26Y or C80I substitutions in AtFKBP12 with other aa changes within the rapamycin-binding domain to make AtFKBP12 more yeast-, human-, or Physcomitrella-like. Or to check whether the introduction of these substitutions in other plant FKBP12 sequences could enhance their affinity to rapamycin. In contrast to Chlamydomonas FKBP12, which has no cysteines in its protein sequence, FKBP12 proteins from *Cyanidioschyzon merolae* and *Auxenochlorella pyrenoidosa* have one cysteine homologous to C80 in land plant FKBP12, and additional cysteines in their extra regions at their N-terminus. These extra regions are believed to prohibit rapamycin binding (Zhu et al. 2022). Therefore, one could test if the deletion of these extra regions could make these two algal species sensitive to the drug. It would also be interesting to test whether other bryophyte species (mosses, liverworts, hornworts) are sensitive to rapamycin. These non-vascular plants express FKBP12 proteins. Intriguingly, the FKBP12 protein sequences of the three mosses *Takakia lepidozioides*, *Calohypnum plumiforme*, and *Sphagnum magellanicum* contain a glutamine at position 58, equivalent to the one in ScFKBP12. Since we showed that introducing this glutamine in PpFKBP12 enhances its affinity to rapamycin, it is reasonable to consider that FKBP12 proteins from these species may have a higher affinity to the inhibitor than the native Physcomitrella FKBP12 has.

The proteins TOR, LST8, and RAPTOR are the main subunits of TORC1 (Loewith et al. 2002). In Chlamydomonas and Arabidopsis, TOR interacts with LST8 *in vitro* and *in vivo* (Díaz-Troya et al. 2008; Moreau et al. 2012). Here, we show that Physcomitrella LST8 substitutes ScLST8 to allow yeast growth, indicating that PpLST8 can be integrated in the yeast TOR complex. Further biochemical experiments will be needed to verify their interaction in Physcomitrella. The line ESPB003 that overexpresses *PpFKBP12* is suited for such an analysis. In fact, in this line, the PpFKBP12 protein is fused at its C-terminus to cMyc, which did not impair its function and could be used to detect the protein in Western blots. Since rapamycin bridges FKBP12 and TOR together, we plan to use this condition to immunoprecipitate PpFKBP12-cMyc using anti-cMyc magnetic beads, and subsequently detect the interacting proteins via Western-blot or LC-MS in forthcoming experiments. However, it should be considered performing this immunoprecipitation *in vitro* and in the presence of a cross-linker to facilitate protein interactions, as described for Chlamydomonas (Díaz-Troya et al. 2008).

In Arabidopsis, TOR plays an essential role throughout all stages of development (Liu et al. 2025). Our results with rapamycin and AZD8055 reveal that Physcomitrella TOR not only controls protonema growth but is also required for developmental progression to gametophores. The Physcomitrella life cycle starts with spores that germinate into protonema, composed of chloronema and caulonema cells. From the protonema, buds develop into shoot-like gametophores bearing male and female sex organs and having root-like structures, called rhizoids. After fertilization, the zygote grows into a sporophyte consisting of a foot, stalk, and capsule, wherein meiosis produces new haploid spores to complete the life cycle (Lueth and Reski 2023). It would be interesting to investigate whether TOR also regulates other developmental stages of Physcomitrella, beyond protonema growth and gametophore formation, especially sexual reproduction (Lüth et al. 2023). The induction by estradiol of the RNAi systems to target PpTORC1 subunit transcripts also affected gametophore formation. However, we had anticipated a more pronounced phenotype, based on observations in Arabidopsis, where a similar system was used (Xiong and Sheen 2012). To our knowledge, the estradiol-inducible RNAi system to downregulate essential genes in Physcomitrella has been predominantly assessed in terms of cell division and viability (Kubo et al. 2013; Kubo et al. 2019). Therefore, comparative data are lacking to place our results into context.

In vascular plants, TOR activity is modulated by diverse environmental cues such as nutrients, energy, light, hormones, biotic and abiotic stresses. This is mediated through well characterized upstream regulators as sensors of these variations (Saliba et al. 2021; Primo et al. 2022; Liu et al. 2025). The downstream effectors through which TOR transduces these signals are also well defined (Liu et al. 2025). Such knowledge was missing for non-vascular plants. In the present study we identified Physcomitrella S6K and RPS6, and showed via LC-MS/MS that TOR-inhibition by the AZD8055 impacts PpRPS6 phosphorylation on Ser241. Given that the Physcomitrella genome is sequenced and well annotated (Lang et al. 2018), identification of additional TOR pathway components should be feasible. Future large-scale phosphoproteomic or other omics studies employing TOR inhibitors, together with genetic and biochemical approaches, will broaden our understanding of the various components of the TOR signalling pathway in this species. The identification of PpRPS6 phosphorylation at Ser241 in a TOR-dependent manner provides a valuable molecular marker that could be used to develop a sensitive Western blot assay of TOR activity using antibodies raised against phosphorylated Ser241 (Dobrenel et al. 2016). Once established, this assay could be used to determine whether Physcomitrella TOR activity responds to the environmental cues described above. As Physcomitrella can be grown in large quantities in bioreactors under highly standardized conditions (Decker and Reski 2004), testing the effects of different environmental parameters is quite straightforward.

## Conclusion

This study closes the knowledge gap about TOR signalling in non-vascular plants, such as mosses. We found a remarkable evolutionary conservation of key components, as well as specific differences, such as the expansion of the *RAPTOR* genes in Physcomitrella and other mosses, compared to Arabidopsis. Such a gene expansion might suggest neofunctionalization of their proteins and indicates differences in signal integration for environmental cues between vascular and non-vascular plants, especially as RAPTOR proteins are regulatory proteins of the TOR complex. Based on sequence analyses, we suggest that other moss species, such as the living fossil *Takakia* may have an enhanced sensitivity to rapamycin, underscoring the necessity for analysing a phylogenetically diverse set of plant species. The tools established here will help to further dissect TOR signalling in Physcomitrella in order to elucidate signalling integration via TORC1 in plants.

## Materials and methods

### Plant material, culture conditions, and dry weight measurement

Physcomitrella WT (new species name: *Physcomitrium patens* (Hedw.) Mitt.; Medina et al. 2019), ecotype “Gransden 2004” (IMSC accession number 41269) and its derivative transgenic lines were cultivated in Knop medium (pH 5.8) supplemented with microelements (ME) as described previously (Frank et al. 2005; Horst et al. 2016). For cultivation on solid medium, 12 g/L agar was added to the Knop ME preparation. Unless noted otherwise, Physcomitrella was grown under standard light conditions (70 µmol photons m^-2^ s^-1^) at 22 °C in a 16 h/8 h light/dark cycle. For stock suspension cultures, moss tissues were grown in 500 mL Erlenmeyer flasks containing 180 mL of Knop ME (pH 5.8). Tissues were disrupted weekly with an ULTRA-TURRAX (IKA, Staufen, Germany) at 18,000 rpm for 60 s to maintain them as protonema. All suspension cultures were subjected to shaking at a speed of 125 rpm. Axenity of the cultures were monitored according to Heck et al. (2021). For dry weight (DW) measurement, 10-25 mL of tissue suspension were vacuum-filtered, dried for 2 h at 105 °C, and then weighed. All measurements were conducted on at least two to three biological replicates. All photos showing Physcomitrella plants on plates or in flasks were taken with either a Canon EOS 400D or Canon EOS 2000D.

### Yeast complementation

Coding sequences were amplified from either yeast genomic DNA or Physcomitrella cDNA and cloned into linearized plasmids (p415 GAL1 or pRL614) using Gibson assembly (Gibson et al. 2009). Point mutations were introduced into *PpFKBP12* (systematic number *Pp3c1_27870*) by site-directed mutagenesis using PCR with synthesized mutagenic primers. The primers were designed to contain the desired nucleotide substitution near the centre of the sequence (all primer sequences are listed in **Supplementary Table S1**). PCR amplification was performed using Phusion™ Plus DNA Polymerase (Thermo Fisher Scientific, Waltham, USA) with plasmids containing *PpFKBP12* (pESPB003) or *PpFKBP12^C26Y-W40D^* (pESPB010) as DNA templates. All generated plasmids were verified by sequencing and are listed in **Table 1**. Yeast strains used in this study were obtained from the laboratory of M.N. Hall (Biozentrum, University of Basel, Switzerland) and are listed in **Table 2**. Plasmids were introduced into yeast cells by transformation using the lithium acetate method (Gietz et al. 1992). Detailed growth conditions for yeast strains are provided in the Figure legends. SD-Leu refers to yeast synthetic dextrose (SD) medium, and SGal-Leu to yeast synthetic galactose (SGal), both of them lacking leucine. Unless otherwise indicated, uracil, tryptophan, and histidine were added to the synthetic media to complement the yeast auxotrophies.

**Table 1.**
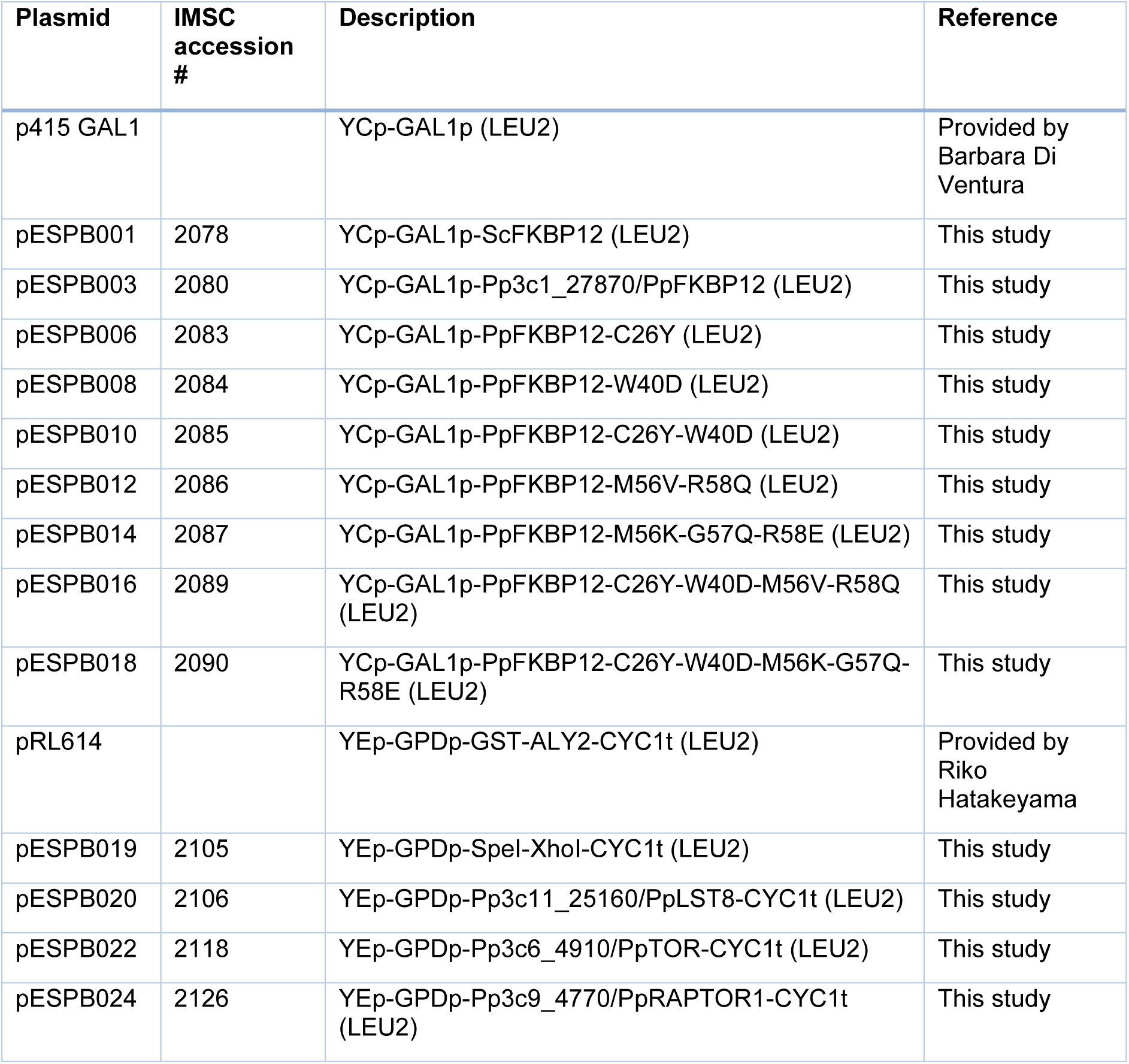
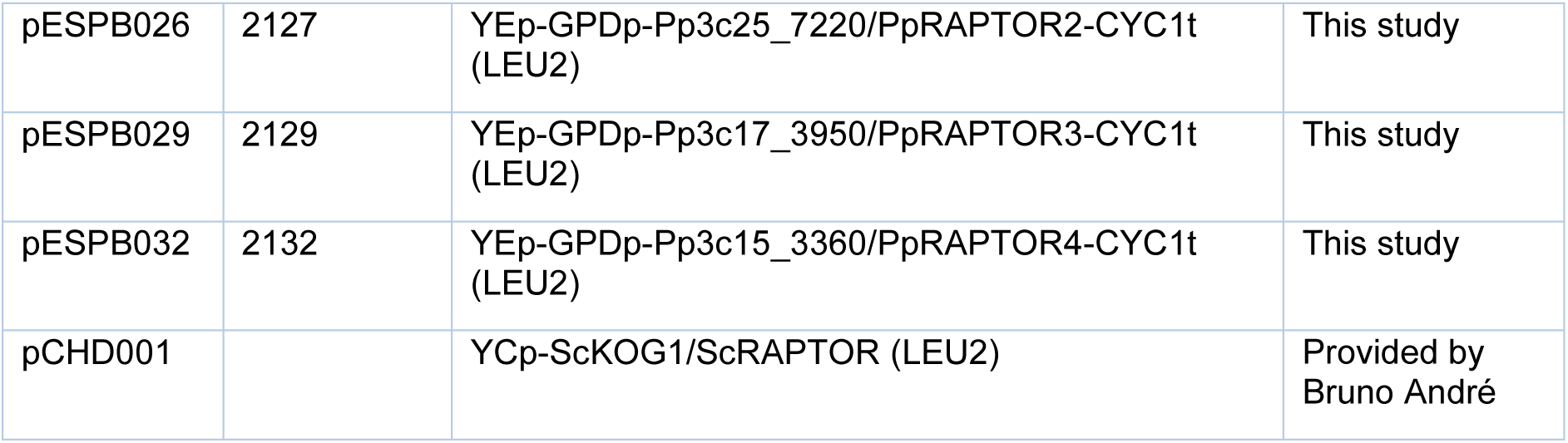
Plasmids used in this study for experiments in yeast.

**Table 2.** Yeast strains used in this study.

| Strain | Genotype |
| --- | --- |
| MH272-3c | MATa *leu2-3,112 ura3-52 rme1 trp1 his3* |
| MH274-1a | MATa *fpr1/fkbp12::URA3 leu2-3,112 ura3-52 rme1 trp1 his3* |
| TB50a | MATa *leu2-3,112 ura3-52 rme1 trp1 his3* |
| SW137-4b | MATa *[HIS3MX6]-GAL1-pTOR2 tor1::[kanMX4] LST8-13myc-[kanMX4] leu2-3,112 ura3-52 rme1 trp1 his3* |
| RL57-2d | MATa *[kanMX4]-GAL1p-LST8 leu2-3,112 ura3-52 rme1 trp1 his3* |
| RL93a | MATa *[kanMX4]-GAL1p-KOG1/RAPTOR leu2-3,112 ura3-52 rme1 trp1 his3* |

### *FKBP12* overexpression lines

DNA constructs for the targeted integration into the Physcomitrella genome of the different *FKBP12* variants fused to *cMyc* were cloned via Gibson assembly. All parts were compiled into a vector backbone containing left (bL) and right (bR) borders of the Physcomitrella intergenic 1 (PIG1) locus, a neutral integration site in the genome (Okano et al. 2009). For each construct, the assembly resulted in a vector that can be linearized via enzymatic digestion to release the following DNA cassette: *PIG1bR-CaMV 35Sp-FKBP12-cMyc-CaMV 35St-PIG1bL*. Coding sequences (CDS) of *ScFKBP12*, *PpFKBP12* and *PpFKBP12-C26Y* were amplified from plasmids pESPB001, pESPB003, and pESPB006, respectively. The CDS of *HsFKBP12* was amplified from *pDONR223-HsFKBP1A/HsFKBP12 SpectR* (provided by Ekkehard Schulze). Transgenic lines were created via highly efficient homologous recombination (Wiedemann et al. 2018) in transformed WT protoplasts (Hohe and Reski 2002; Hohe et al. 2004). The linear DNA cassette and a co-transfection vector conferring resistance on hygromycin B were used for transfection as described before. All DNA materials used for transfection were purified and sterilized via ethanol precipitation before usage (Sambrook and Russell 2006).

Screening of plants surviving hygromycin B selection was done by PCR to validate correct integration of the linear DNA cassette at the PIG1 locus. For this purpose, we used two different primer pairs: one pair (5_Up_173long_PIG1-5 and 3_103_35S-T(inv)) binds to a region in the genome upstream of PIG1bL and in the CaMV 35S terminator, while the other (5_52_35S-P(inv) and 3_Down_81_PIG1-3) binds in the CaMV 35S promoter and a region in the genome downstream of PIG1bR. Because untargeted integrations can still take place, transgene copy numbers were quantified via qPCR as described in Noy-Malka et al. (2014). Genomic DNA was isolated from 100 mg fresh weight (FW) of frozen protonema tissue using the GeneJET Plant Genomic DNA Purification Kit (Thermo Fisher Scientific), following the manufacturer’s protocol. DNA samples were normalized to a concentration of 3 ng/µL prior to qPCR analysis. Reactions were set up using primers targeting PIG1bL (5-qPCR-PIG1-5HR-a and 3-qPCR-PIG1-5HR-a) and PIG1bR (5-qPCR-PIG1-3HR-b and 3-qPCR-PIG1-3HR-b), as well as primers specific for the CaMV 35S promoter (5-qPCR-35Sp(mAV)-c and 3-qPCR-35Sp(mAV)-c). To serve as an internal reference, primers amplifying the single-copy gene *CLF* (*Pp3c22_2940V3*) were included. All primer sequences are listed in **Supplementary Table S2**. WT plants and an in-house reference line *(ΔPpAARE3 #17*; Hoernstein et al. 2023) with a confirmed single integration of a DNA cassette were used as controls. All qPCR reactions were performed using 2x SensiFAST Mix (Bioline, London, UK) on a Lightcycler 480 II system (Roche, Basel, Switzerland). Raw qPCR data are listed in **Supplementary Table S5**.

### Inducible RNA interference lines

To generate the estradiol-inducible *tor*, *lst8*, and *raptors* RNA interference (RNAi) lines of Physcomitrella, we followed the method of Miki et al. (2016) with minor modifications. For the construction of the different RNAi systems destined for a targeted integration at the PIG1 locus, we used the pGG624 plasmid (Miki et al. 2016). For each *TORC* gene subunit, two distinct regions within the transcript sequence were selected as RNAi targets. This approach reduces the likelihood of off-target effects, as consistent phenotypes across both targets strengthen evidence for a gene-specific knockdown. For *PpTOR*, these regions were of 500 and 510 bp length, and covered regions 561-1060 and 2496-3005 in the CDS, respectively. For *PpLST8*, the first target sequence was 500 bp long and covered region 213-712 in the CDS. The second was 435 bp long and was located in the 5’UTR prior to the start codon. To downregulate the expression of the four *PpRAPTOR* paralogs with a single RNAi construct, regions in their CDS that are highly similar were selected. This strategy was described before and was proven successful (Miki et al. 2016). Thus, the first *Ppraptor* RNAi construct contained region 1656-2166 (511 bp) from the CDS of *PpRAPTOR1*. The second contained region 3662-4103 (442 bp) from *PpRAPTOR2*. These regions were 74-85% similar between the four paralogs. All primers used for cloning are listed in **Supplementary Table S3**. β-Estradiol (E8875, Sigma-Aldrich, St. Louis, USA) was used to induce gene silencing.

### Crude extracts and Western blotting

Crude extracts for Western blot analysis were prepared from 100 mg of FW protonema, which were vacuum-filtered, collected in 2 mL Safe-Lock tubes, and frozen in liquid nitrogen. Tissues were homogenized and disrupted using two beads (glass and metal) in a bead mill (MM 400; Retsch, Haan, Germany) for 3 min at 30 Hz. To the ground plant material, 500 µL of extraction buffer (25 mM HEPES-KOH pH 7.5, 100 mM NaCl, 0.2 mM DTT, 2 mM EDTA, 0.5% Triton, 1% plant protease inhibitor cocktail P9599 from Sigma Aldrich) were added to each sample, followed by sonication for 15 min using a cold ultrasound bath (Sonorex RK52, Bandelin, Berlin, Germany), and then centrifuged at 20,000 x g for 30 min at 2°C. Collected supernatants were subjected to another round of centrifugation for 5 min, and at the end collected into new tubes. Samples were quantified using the Pierce™ BCA Protein Assay Kit (Thermo Fisher Scientific) according to the manufacturer’s protocol. Before loading on SDS gel, samples were reduced by heating 10 min at 95°C following the addition of 50 mM DTT and 4x Laemmli sample buffer (Bio-Rad Laboratories, Hercules, USA). 30 µg of total protein were loaded on a 12% acrylamide gel (Mini-PROTEAN® TGX™ Stain-Free™ gels, Bio-Rad Laboratories) and run for 90 min at 120 V. Proteins were transferred to a PVDF membrane (Hybond P0.45, Amersham, Cytiva, Marlborough, USA) in a Trans-Blot SD Semi-Dry Electrophoretic Cell (Bio-Rad Laboratories) for 80 min with 1.5 mA/cm^2^. After blocking with 4% ECL™ Blocking Agent (Cytiva, Amersham), the membrane was probed with mouse monoclonal anti-cMyc antibody (dilution 1:5000 in 1x TBST; clone 9E10, Roche). Anti-cMyc antibody was detected with a horseradish-peroxidase-conjugated anti-mouse IgG secondary antibody from sheep (Cytiva), followed by enhanced chemiluminescence (dilution 1:5000 in 1x TBST; Amersham ECL Prime Western Blotting Detection Reagent, Cytiva). Western blot signals were visualized using autoradiography films (CL-XPosure™ Film, Thermo Scientific, Cat. No. 34090). Transfer quality and total protein levels were monitored by PageBlue™ (Thermo Fisher Scientific) staining of the SDS gels.

### Chlorophyll content

Chlorophyll was extracted from 20 mg protonema (vacuum-filtered material and frozen in liquid nitrogen). Tissues in a 2 mL Safe-Lock tube were homogenized and disrupted using two beads (glass and metal) in a mixer mill (MM 400, Retsch GmbH) for 3 min at 30 Hz, followed by the addition of 480 µL cold 80% acetone and incubation on ice. After mixing for 5 min at Vmax using an Eppendorf ThermoMixer, samples were centrifuged at 20,000 x g for 10 min at 2°C and 400 µL of supernatants were collected. After an additional round of centrifugation for 5 min, three technical replicates of 100 µL of each supernatant were loaded on a 96-well microplate (Greiner, PS, F-Bottom, Crystal-Clear) and absorbance was read at 645 nm and at 663 nm using a POLARstar Omega or a SPECTROstar^Nano^ microplate reader (BMG LABTECH, Ortenberg, Germany). Total chlorophyll was calculated as chlorophyll *a* + chlorophyll *b* (mg g^-1^ FW) using the following formula as described in Ruibal et al. (2020): Chl *a* (mg·g⁻¹ FW) = [(12.7 × A₆₆₃) - (2.6 × A₆₄₅)] × (mL acetone/mg FW); Chl *b* (mg·g⁻¹ FW) = [(22.9 × A₆₄₅) - (4.68 × A₆₆₃)] × (mL acetone/mg FW).

### Light microscopy

A Zeiss Axioplan 2 optical microscope equipped with an AxioCam MRc5 (Zeiss) was used to visualize protonema tissues. For this purpose, 15 µL of protonema suspension were placed on a glass slide, which was then covered with a coverslip before visualization at 10x magnification.

### Chlorophyll fluorescence

Protonema suspension cultures at a density of 250 mg/L DW were dark-adapted for 10 min, and chlorophyll fluorescence was measured via pulse-amplitude modulation (PAM) in an imaging PAM instrument (IMAG-MAX/L, Walz, Effeltrich, Germany) equipped with a Maxi-head (IMAG-K7, Walz). PSII maximum quantum yield (F_v_/F_m_) was recorded after F_0_ and F_m_ determination. All experiments on photosynthetic parameters were repeated independently three times.

### Total protein content

Total protein content was determined from 20 mg protonema (vacuum-filtered material and frozen in liquid nitrogen). Tissues in a 2 mL Safe-Lock tube were homogenized and disrupted using two beads (glass and metal) in a mixer mill (MM 400, Retsch GmbH) for 3 min at 30 Hz, followed by addition of 100 µL extraction buffer (50 mM PBS and 1 mM EDTA). Samples were vigorously hand-shaken, centrifuged for 5 sec at 1.5 x g, and vortexed for 10 sec. To enhance extraction, samples were further processed in the mixer mill for 5 min at 20 Hz. This was followed by sonication in a cold ultrasonic bath (Sonorex RK52, Bandelin) for 15 min. The lysates were centrifuged at 20,000 × g for 30 min at 2°C. The supernatants were carefully collected and subjected to a second centrifugation step (5 min), after which the supernatants were transferred to new tubes. Total protein content was determined from 1:4 diluted samples in extraction buffer using the Pierce™ BCA Protein Assay Kit (Thermo Fisher Scientific) according to the manufacturer’s instructions.

### Flow cytometry

Flow cytometry analysis was conducted as previously described (Heck et al. 2021) with slight modifications. Protonema tissues were chopped with a razor blade in a small petri dish (6 cm diameter) containing 0.5 mL of DAPI-buffer (0.01 mg/L 4′,6-Diamidin-2-phenylindol (DAPI), 1.07 g/L MgCl2 ×6 H2O, 5 g/L NaCl, 21.11 g/L Tris, 0.1% Triton, pH 7). Subsequently, 1.5 mL DAPI-buffer was added to the preparation, which was then filtered through a sieve of 30 µm pore size. The filtrate was analysed with a Partec CyFlow® Space flow cytometer (Sysmex Partec, Görlitz, Germany) equipped with a 365 nm UV-LED.

### AZD treatment for phospho-proteomic analysis

Moss suspension culture normalised to 180 mg/L was divided into 50 mL Erlenmeyer flasks and cultivated for 24 h under standard growth conditions. Subsequently, they were incubated for 1 h with either 10 µM AZD8055 (AZD), or an equivalent amount of DMSO as a control. Treated tissue was harvested by vacuum-filtration, transferred to a 2 mL SafeLock reaction tube (Eppendorf) containing one 3 mm tungsten carbide bead (Qiagen) and one glass bead. Samples were snap-frozen in liquid nitrogen and ground to fine powder at 28 Hz for 2 min in a homogenizer (MM 400, Retsch). Lysis buffer according to Lang et al. (2011) containing 7.5 M urea, 2.5 M thiourea, 12.5% glycerol, 62.5 mM Tris-HCl, 2.5% (w/v) n-octyl glucoside (octyl β-D-glucopyranoside), 1.25 mM protease inhibitor P2714 (Sigma-Aldrich) was supplemented with PhosStop phosphatase inhibitor (Roche) and added to the ground tissue. Samples were vortexed until dissolved, incubated for 15 min in an ultrasound bath, and centrifuged at 20,000 x g and 16 °C for 30 min. Finally, the supernatant containing the extracted protein was transferred to a fresh reaction tube. The protein concentration was determined using the Quick Start Bradford Protein Assay (Bio-Rad) according to the manufacturer’s instructions.

### SP3 clean up, tryptic digest and phospho-peptide enrichment

The extracted proteins were further processed using an adapted single-pot, solid-phase-enhanced sample-preparation (SP3) method (Hughes et al. 2019). Briefly, samples (200 µg input) were reduced with 2 mM tris(2-carboxyethyl) phosphine (TCEP) for 30 min at 37 °C, then alkylated with 50 mM chloroacetamide (CAA) for 30 min in the dark, and finally, the reaction was quenched with 50 mM DTT for 20 min at RT. Sera-Mag SpeedBead magnetic carboxylate-modified particles (Cytiva) were washed twice with 10x volume of ELGA H_2_O, then added in a 1:10 bead to proteome ratio to the samples and incubated for 20 min at RT in a high organic solution of 80% EtOH to allow the proteins to bind to the beads. The beads were bound using a magnetic rack and washed twice with 10x volume of 90% acetonitrile (ACN). To resuspend the beads in between washing steps, the tubes were placed in an ultrasonic water bath. After washing, the remaining liquid was removed, and the beads were resuspended in 100 mM ammonium bicarbonate (AMBIC). Trypsin was added in a 1:100 protease to proteome ratio and incubated at 37 °C for 16 h and shaken at 1,200 rpm. Digested peptides were desalted using µSpin C18-disc cartridges (Affinisep) following the manufacturer’s instructions. After elution in 50 % ACN and 0.5 % acetic acid (HAC) the peptides were dried using a Speedvac concentrator (Eppendorf).

Peptides were reconstituted in 1 µL binding buffer (80% ACN, 1% trifluoracetic acid (TFA), 3% (w/v) glycolic acid (GA)) per 1 µg dried peptide. The magnetic Ti-NTA beads (Cube Biotech, Monheim, Germany) were washed three times in 10 vol of binding buffer, resuspended in binding buffer, then incubated for 30 min at 25 °C and 1200 rpm with the peptides in a 10:1 (w/w) bead to peptide ratio. After binding, the beads were washed twice in wash buffer 1 (80% ACN, 1% TFA), and once in wash buffer 2 (10 % ACN, 0.2% TFA), in between wash steps, the beads were placed in an ultrasonic water bath and incubated for 2 min at 25 °C and 1200 rpm. Finally, the peptides were incubated for 30 min at 25 °C and 1200 rpm in elution buffer (1% ammonium hydroxide (NH_4_OH)), then the supernatant containing the eluted peptides was transferred to a fresh reaction tube and acidified to pH <3 using formic acid.

Acidified peptides were desalted using self-packed SDB-RPS Stop and Go Extraction tips (StageTips) (Rappsilber et al. 2007) composed of three layers of 1.0 × 1.0 mm (AttractSPE Disks Bio RPS, Affinisep, Le Houlme, France). The StageTips were equilibrated sequentially with 100% methanol, 80% acetonitrile containing 0.1% FA, and twice with 0.1% FA. Acidified peptide solutions were loaded onto the StageTips and washed with 0.1% FA, 80% acetonitrile with 0.1% FA, and 80% methanol with 0.5% FA. The eluates were dried using a vacuum concentrator (Eppendorf) and reconstituted in 0.1% FA prior to LC-MS measurement.

### Measurement on a QExactive Orbitrap

For LC-MS analysis of the phospho-peptide enriched samples, an UltiMate 3000 RSLCnano system was coupled online to a QExactive mass spectrometer (both Thermo). The peptide mixture was washed and preconcentrated on a PepMap C18 trapping column (Thermo) with a flow rate of 30 µL/min and peptides were separated on a μPAC C18 pillar array column (200 cm bed length, Thermo) using a binary buffer system consisting of A (0.1% formic acid) and B (86% acetonitrile, 0.1% formic acid). Peptides were eluted applying a linear gradient from 5% to 50% B in 45 min and 50 to 99% B in 2 min at a flow rate of 1 µL/min. For electrospray ionization of peptides, a Nanospray Flex ion source (Thermo Fisher Scientific) with a liquid Junction PST-HV-NFU (MS Wil, Aarle-Rixtel, The Netherlands) and a fused silica emitter (20 µm inner diameter, 360 µm outer diameter, MicroOmics Technologies LLC) applying a source voltage of +1.800 V and an ion transfer tube temperature at 250°C was used. Cycles of data-independent acquisition (DIA) consisted of: one overview spectrum (RF lens of 50%, normalized AGC target of 3e6, maximum injection time of 60 ms, m/z range of 385 to 1.050, resolution of 70,000 at 200m/z, profile mode) followed by MS^2^ fragment spectra (RF lens of 50%, normalized AGC target of 1e6, maximum injection time of 60 ms) generated sequentially by higher-energy collision-induced dissociation (HCD) at a normalized energy of 26% in a precursor m/z range from 400 to 1,000 in 50 windows of 12.5 m/z isolation width overlapping by 0.25 m/z and recorded at a resolution of 35,000 in profile mode.

### Database search with *FragPipe*

Raw mass spectrometry files of phospho-peptide enriched samples were analysed using *FragPipe* v22.0 configured for data independent acquisition (DIA). Peptide identification was performed using *MSFragger* (Kong et al. 2017) against a *Physcomitrium paten*s protein database (48,582 entries; Lang et al. 2018) containing both target and decoy sequences for FDR estimation. The search was carried out with a precursor mass tolerance of −20 to +20 ppm, enzyme specificity set to trypsin, a peptide mass range of 500 to 5,000 Da, and a minimum peptide length of 7 and a maximum of 40 amino acids. Variable modifications included phosphorylation of serine, threonine, and tyrosine (S/T/Y), oxidation of methionine (M), deamidation of asparagine and glutamine (N/Q), pyro-glutamate conversion from N-terminal glutamine, glutamic acid, and cysteine (Q/E/C), as well as hydroxylation of proline (P). Validation of peptide-spectrum matches, and phospho-site assignments were performed using *MSBooster* (Yang et al. 2023), Percolator (Käll et al. 2007; The et al. 2017), and PTMProphet (Shteynberg et al. 2019). Quantification was conducted using *DIA-NN* (Demichev et al. 2020), applying a protein-level false discovery rate (FDR) threshold of 0.01.

### Phospho-peptide data analysis with R

Phospho-peptide data analysis was performed in *R* (v4.4.3; R Core Team 2025), using a minimum site localization probability of 0.75 for serine, threonine, and tyrosine (S/T/Y) residues. Statistical significance of differences between conditions was assessed using a Welch test.

### Protein sequence alignment, phylogenetic inference and functional domain analysis

FKBP12 protein sequences from *A. thaliana* (AT5G64350.1) and *P. patens* (Pp3c1_27870V3.1) were complemented by homologs from *C. reinhardtii*, *Homo sapiens*, *O. sativa*, *S. cerevisiae*, *Schizosaccharomyces pombe* (Crespo et al. 2005), *A. pyrenoidosa* (Zhu et al. 2022), *C. merolae* (Imamura et al. 2013), *S. lycopersicum* (Xiong et al. 2016), and *V. faba* (Xu et al. 1998). Based on this set we conducted proteome-wide sequence searches with Diamond (v2.1.10.164; Buchfink et al. 2021) against *Anthoceros angustus* (Zhang et al. 2020), *Calohypnum plumiforme* (Mao et al. 2020), *Ceratopteris richardii* (Marchant et al. 2022), *Marchantia polymorpha* (Bowman et al. 2017), *Populus trichocarpa* (Tuskan et al. 2006), *Selaginella moellendorffii* (Banks et al. 2011), *Sphagnum magellanicum* (Healey et al. 2023), *Takakia lepidozioides* (Hu et al. 2023), and *Vitis vinifera* (Jaillon et al. 2007). InterProScan (v5.71-102.0; Jones et al. 2014) was used to infer functional domain annotation. Initial hits were filtered for assignment to the Panther (Thomas et al. 2022) family “PEPTIDYL-PROLYL CIS-TRANS ISOMERASE” (PTHR10516), manually curated and aligned with Mafft (v7.490; Katoh and Standley 2013) in “globalpair” mode.

We performed Diamond-based sequence searches with RAPTOR homologs from *A. thaliana* and *P. patens* against the proteomes of *A. angustus*, *Ceratodon purpureus* (Carey et al. 2021), *C. richardii*, *Funaria hygrometrica* (Kirbis et al. 2025), *Klebsormidium nitens* (Hori et al. 2014)*, Mesotaenium endlicherianum* (Cheng et al. 2019)*, M. polymorpha*, *O. sativa*, *P. trichocarpa*, *S. moellendorffii*, *T. lepidozioides*, and *V. vinifera*. Initial hits were annotated with InterProScan, filtered for assignment to Panther family “REGULATORY-ASSOCIATED PROTEIN OF MTOR” (PTHR12848) and aligned with Mafft in “localpair” mode. For phylogenetic reconstruction we identified ‘JTT+I+G4+F’ as best-fit model with ModelTest-NG (v0.1.7; Darriba et al. 2020) and calculated a maximum likelihood tree with 1000 bootstrap replicates using RAxML-NG (v1.2.2; Kozlov et al. 2019). The tree was visualized and rooted with iTOL (v7; Letunic and Bork 2024).

We inferred functional domains for TOR, RAPTOR and LST8 homologs from *A. thaliana*, *O. sativa*, *P. patens*, and *S. cerevisiae* (Liu et al. 2025) using InterProScan (v5.71-102.0; Jones et al. 2014). Pfam (Paysan-Lafosse et al. 2024) domains “FKBP12-rapamycin binding domain” (PF08771), “FAT domain” (PF02259), “Phosphatidylinositol 3- and 4-kinase” (PF00454), “FATC domain” (PF02260), and “Raptor N-terminal CASPase like domain” (PF14538) were relabelled as “FRB”, “FAT”, “KD”, “FATC”, and “RNC”, respectively. Superfamily (Pandurangan et al. 2018) annotations “ARM repeat” (SSF48371), and “WD40 repeat-like” (SSF50978) were relabelled as “HEAT”, and “WD40”, respectively, and consecutive “WD40” domains were concatenated for sequence comparison and visualization. Pairwise sequence identity was calculated using R (v4.4.3; R Core Team 2025) and the pwalign package (v1.2.0; Aboyoun et al. 2025), and protein domain plots were generated using the seqvisr (v0.2.7; vragh 2022) package.

### Statistical analysis

Graphs and statistical analyses were performed with the GraphPad Prism software version 10.0 for Windows (GraphPad software, San Diego, California, USA). Statistical differences were analysed with one-way Anova with subsequent post-hoc test in R. Statistical significance was accepted at *p < 0.05, **p < 0.01, and ***p < 0.001.

## Author contributions

E.S. designed research, acquired funding, performed experiments, analysed data, and wrote the manuscript. S.N.W.H., N.v.G, A.S., and K.M.V.H. designed research, performed experiments, analysed data, and helped writing the manuscript. J.P., E.L.D., and P.F.H. analysed data and helped writing the manuscript. H.T.S. supervised research. R.R. designed and supervised research, analysed data, acquired funding, and wrote the manuscript. All authors read and approved the final version of the manuscript.

## Data availability

The WT strain as well as the transgenic lines generated in this study are accessible via the International Moss Stock Center (IMSC, www.moss-stock-center.org). Their IMSC accession numbers are listed in **Supplementary Table S4**. The mass spectrometry proteomics data have been deposited at the ProteomeXchange Consortium via the PRIDE partner repository (Deutsch et al. 2023; Perez-Riverol et al. 2025) with the dataset identifier PXD069889.

## Acknowledgements

We are grateful to Michael N. Hall, Barbara Di Ventura, Bruno André, Claudio De Virgilio, Riko Hatakeyama, Elena Kozgunova, Sjon Hartman, and Ekkehard Schulze for strains and plasmids. We thank Sarah Courbier for including our samples in some of her TOR activity assays. We also thank Agnes Novakovic and Richard Haas for technical assistance, and Anne Katrin Prowse for language editing.

## Supplementary information

**Supplementary Figure S1.**
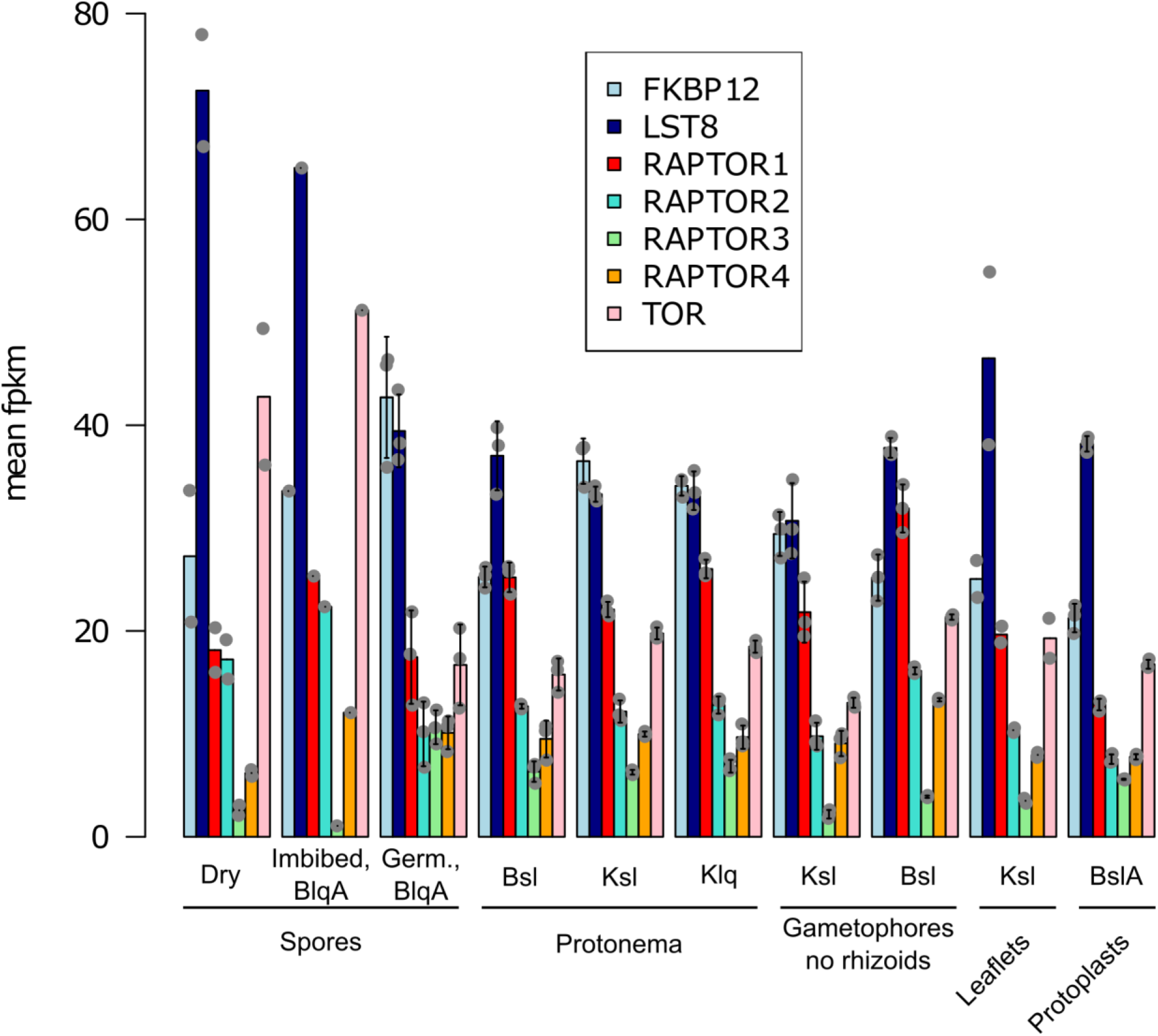
Expression levels of TORC1 component genes in Physcomitrella at different culture conditions and developmental stages. Expression data were retrieved from *PEATmoss* (https://peatmoss.plantcode.cup.uni-freiburg.de/expression_viewer/input). Values represent mean FPKM values (Fragments Per Kilobase Million) and standard deviation. Data points represent biological replicates. Abbreviations: B = BCD medium; lq = liquid; A = ammonium tartrate; sl = solid; K = Knop medium; Germ. = germinating.

**Supplementary Figure S2.**
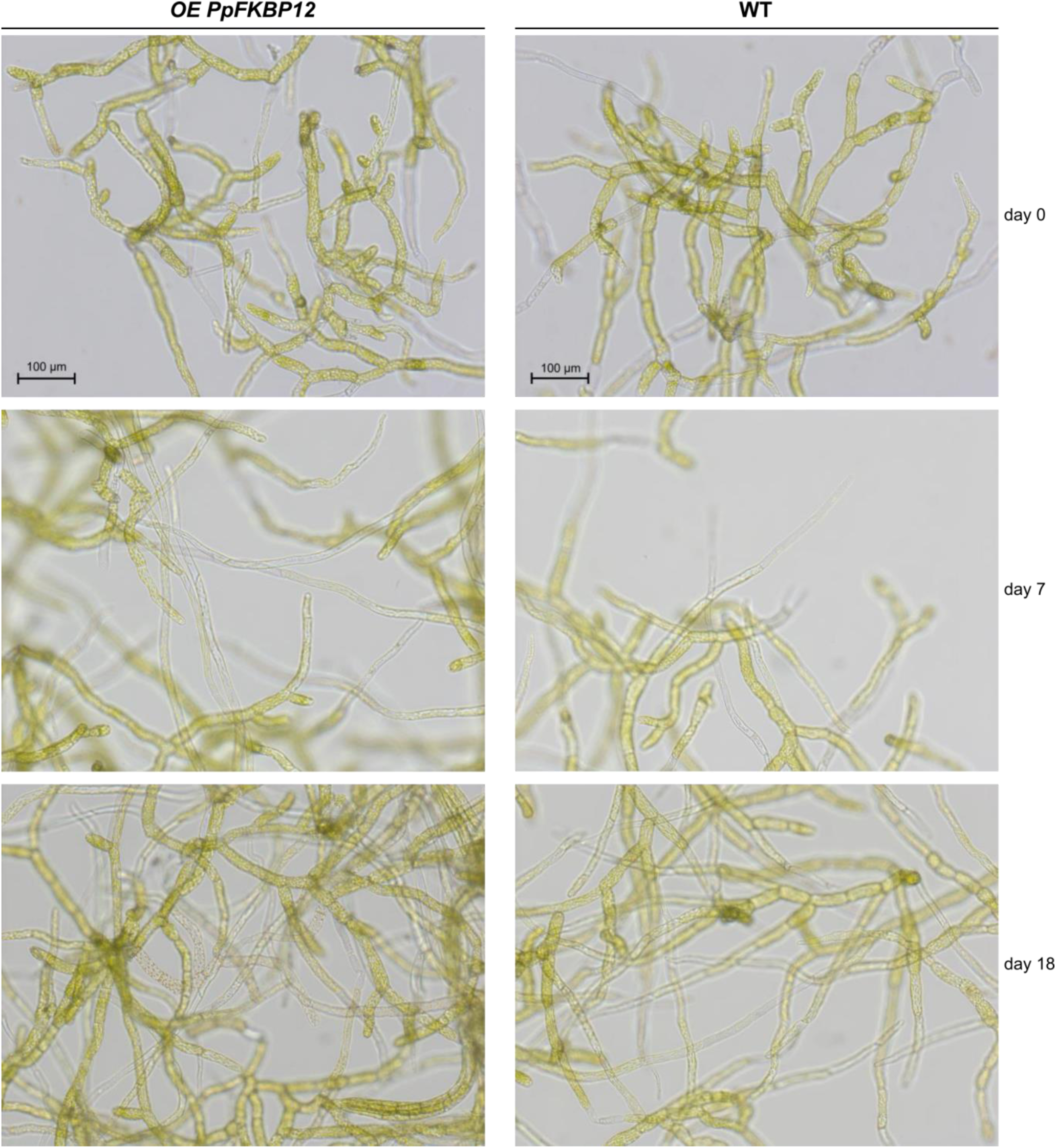
Caulonema frequency increased with incubation time. Light microscopy images of protonemata in suspension cultures treated with DMSO 0.01 % and incubated for 7 and 18 days. Culture conditions are similar to those in Figure 1.

**Supplementary Figure S3.**
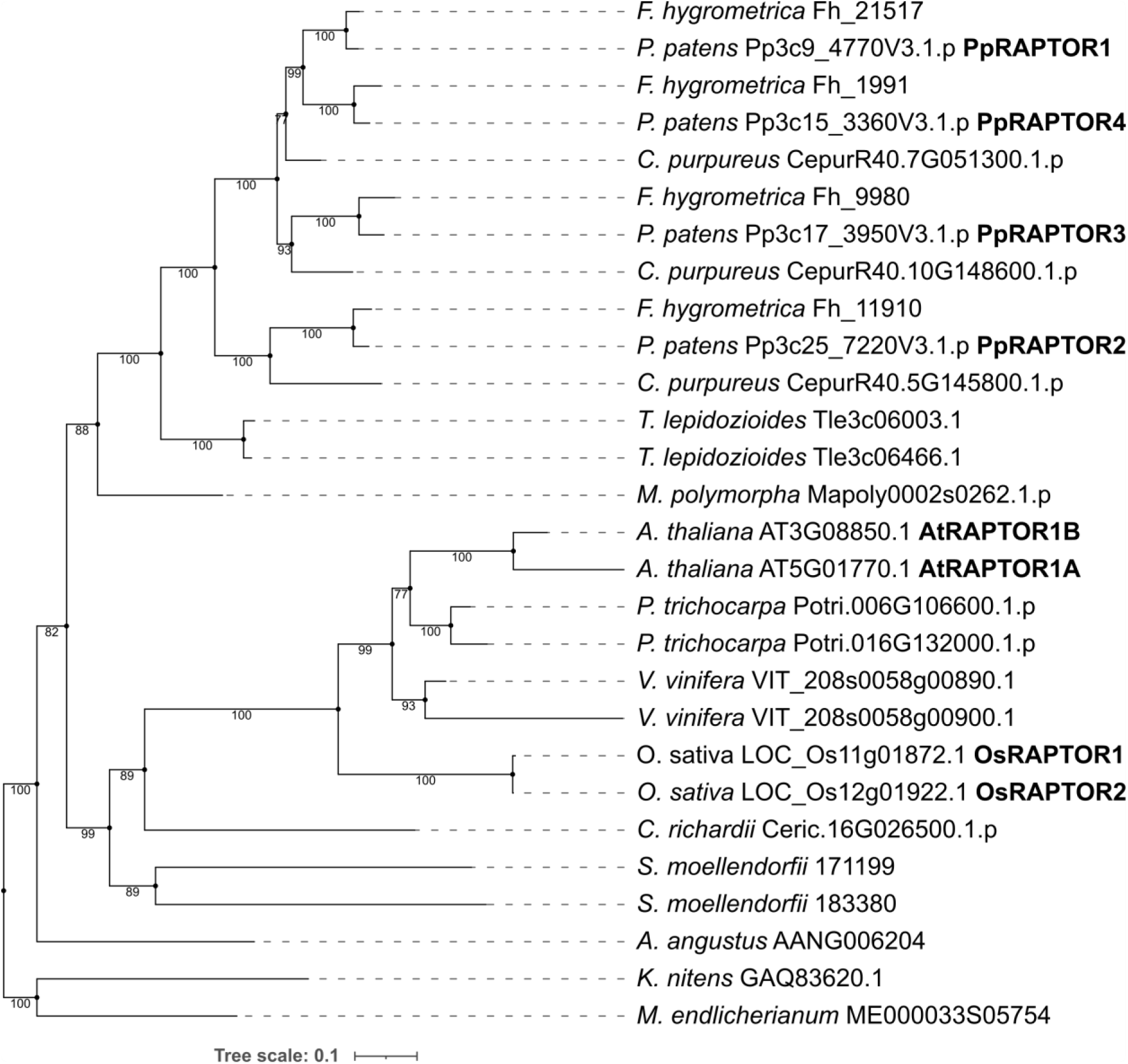
Phylogeny of the plant RAPTOR family. Maximum-likelihood tree based on a multiple sequence alignment of RAPTOR proteins, rooted at the split between streptophytic algae and land plants. Common names of RAPTOR isoforms are written in bold where available. Numeric values at nodes indicate percentage support based on 1000 bootstrap trees.

**Supplementary Figure S4.**
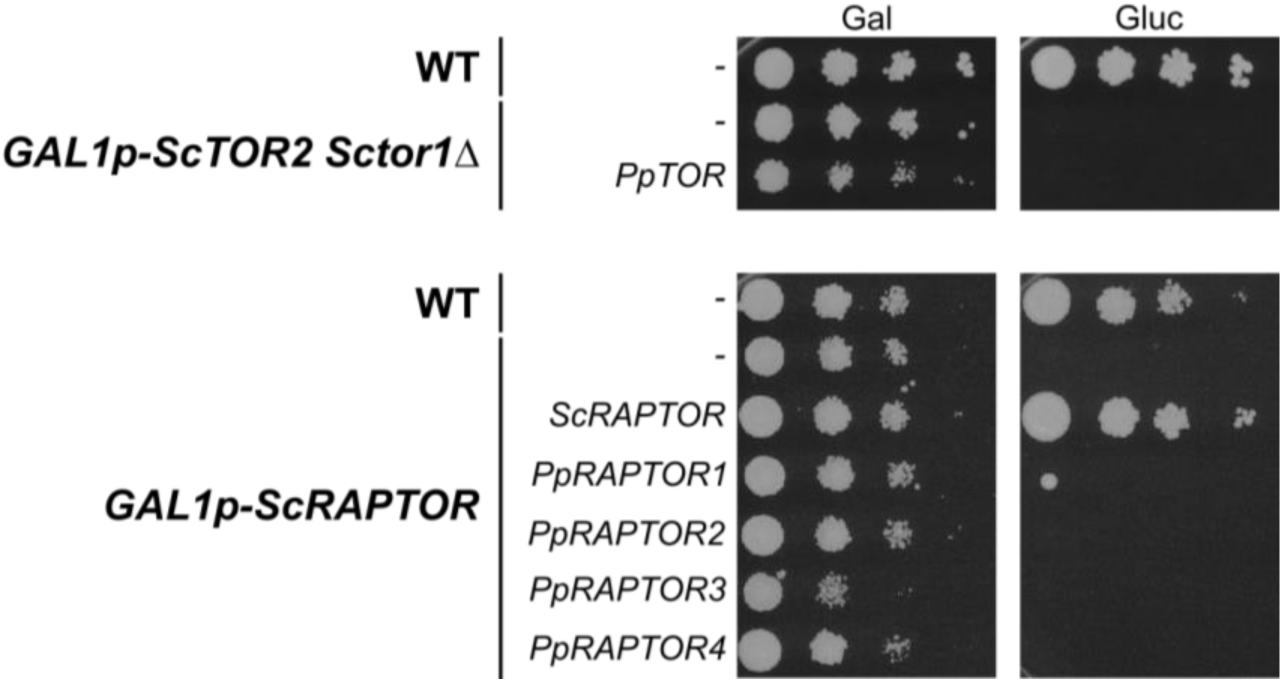
Complementation of the yeast *tor* and *raptor* mutants with their Physcomitrella homologs. **A** Wild-type TB50a yeast strain and mutant SW137-4b expressing *ScTOR2* from the galactose-inducible GAL1 promoter were initially grown as suspensions in rich medium containing galactose as a carbon source (YPGal). Strains were transformed with pESPB019 (empty vector) or pESPB022 (pESPB019-derived vector expressing *PpTOR*) and selected on SGal-Leu plates. Selected transformants were grown for 6 h at 30°C in SD-Leu suspension cultures. Cultures were normalized to an OD (660 nm) of 0.2, subjected to 10-fold serial dilutions, and 15 µl of each dilution were spotted onto SGal-Leu or SD-Leu plates, which were incubated at 30°C for 5 days. Plates were scanned upside down with the lid open on a Canon CanoScan 9000F scanner. **B** Same growth conditions as in panel A. Strain RL93a expressing yeast *RAPTOR* from the GAL1 promoter was transformed with pESPB019 (empty vector), pESPB019-derived vectors each containing one of the Physcomitrella RAPTOR paralogs (pESPB024, pESPB026, pESPB029, and pESPB032), or pCHD001 expressing *ScRAPTOR*.

**Supplementary Figure S5.**
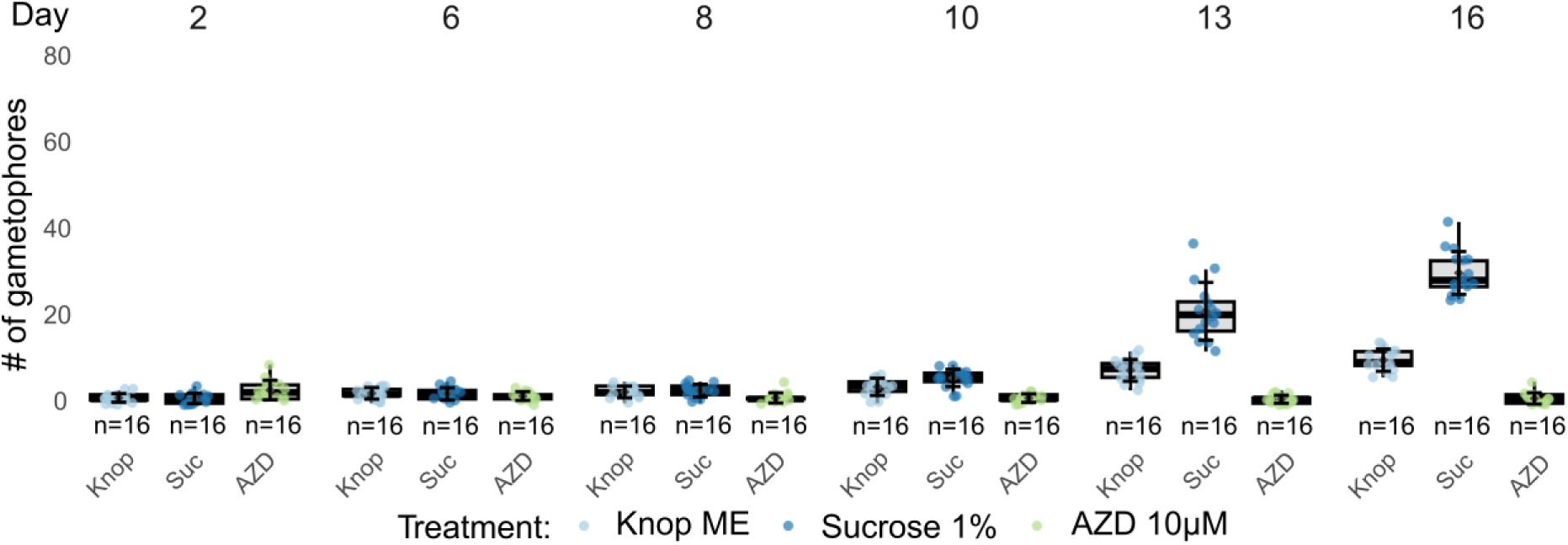
Gametophore counts over time in Physcomitrella WT. Protonema at a density of 440 mg DW/L was spotted on solid medium plates (15 µL) containing either only Knop ME or additionally 1% sucrose or 10 µM AZD8055. Arising gametophores were counted over 16 days. Each data point represents a single spot at which gametophores were counted. n = 16 colonies per treatment.

**Supplementary Figure S6.**
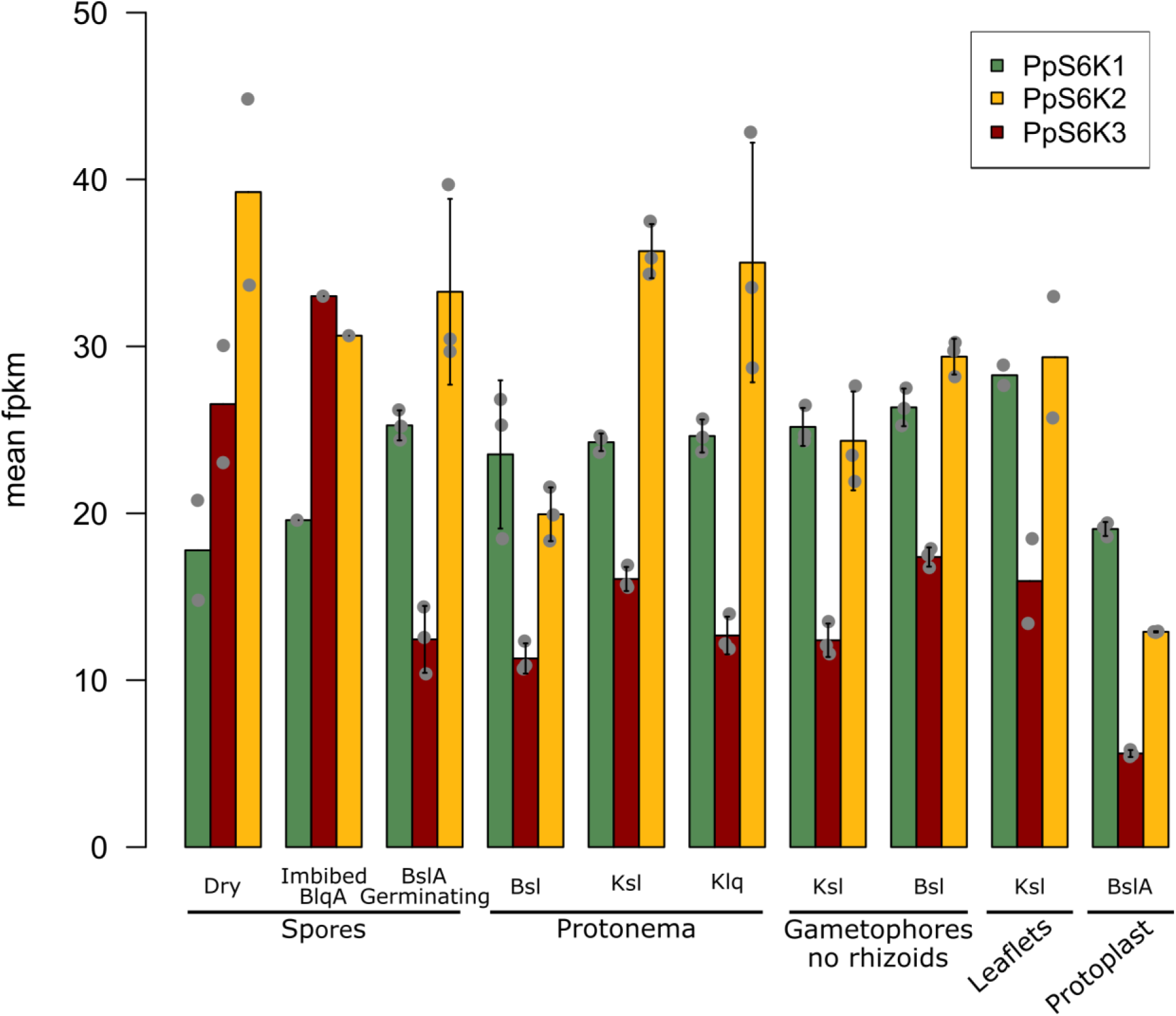
Expression levels of *PpS6K* genes in Physcomitrella at different culture conditions and developmental stages. Expression data were retrieved from *PEATmoss* (https://peatmoss.plantcode.cup.uni-freiburg.de/expression_viewer/input). Values represent mean FPKM values (Fragments Per Kilobase Million) and standard deviation. Data points represent biological replicates. Abbreviations: B = BCD medium; lq = liquid; A = ammonium tartrate; sl = solid; K = Knop medium; Germ. = germinating. PpS6K1: Pp3c12_223 60V3.1; PpS6K2: Pp3c25_14820V3.1; PpS6K3: Pp3c16_24840V3.1

**Supplementary Figure S7.**
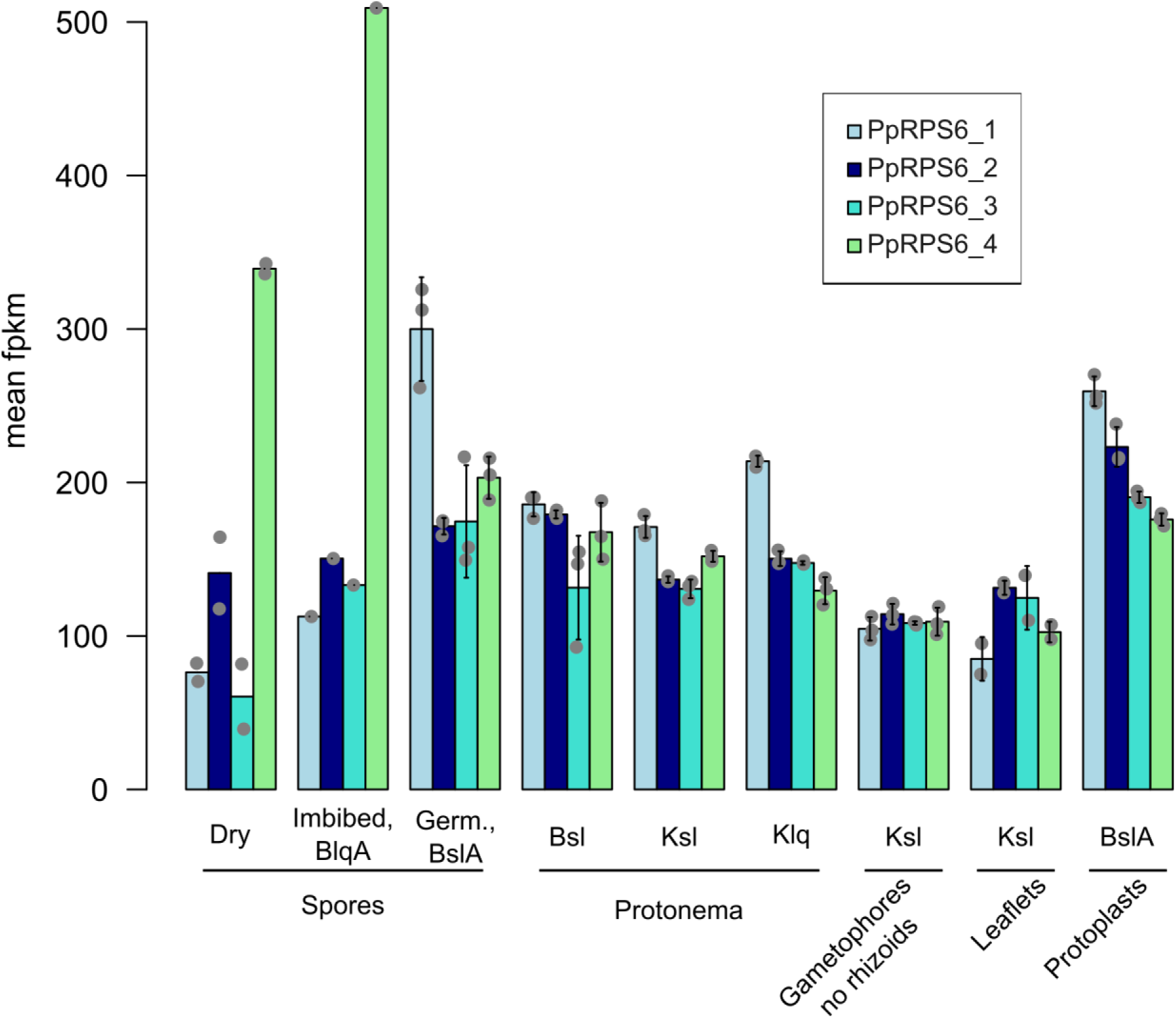
Expression levels of *PpRPS6* genes in Physcomitrella at different culture conditions and developmental stages. Expression data were retrieved from *PEATmoss* (https://peatmoss.plantcode.cup.uni-freiburg.de/expression_viewer/input). Values represent mean FPKM values (Fragments Per Kilobase Million) and standard deviation. Data points represent biological replicates. Abbreviations: B = BCD medium; lq = liquid; A = ammonium tartrate; sl = solid; K = Knop medium; Germ. = germinating. PpRPS6_1: Pp3c11_21120V3.1; PpRPS6_2: Pp3c15_25430V3.1; PpRPS6_3: Pp3c15_25780V3.1; PpRPS6_4: Pp3c7_5780V3.1

**Supplementary Figure S8.**
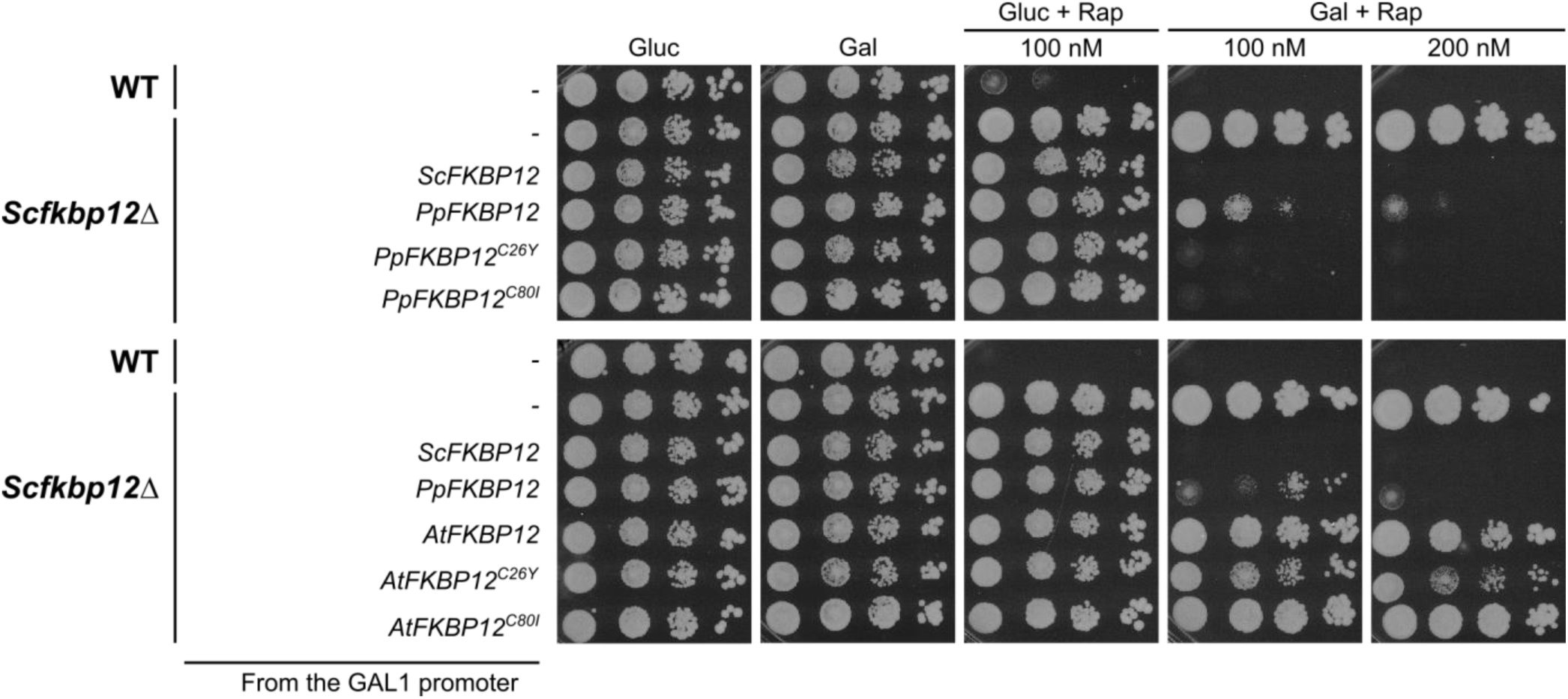
Yeast complementation assay of the *fkbp12Δ* mutant with mutant FKBP12 variants from Physcomitrella and Arabidopsis. Wild-type MH272-3c yeast strain transformed with an empty vector (p415 GAL1), and mutant MH274-1a (*fkbp12Δ*) strain transformed with either p415 GAL1 or p415 GAL1-derived vectors expressing different *FKBP12* variants, were grown in SD-Leu or SGal-Leu suspension cultures. Cultures were normalized to an optical density (OD) at 660 nm of 0.1, subjected to 10-fold serial dilutions, and 15 µL of each dilution were spotted onto SD-Leu or SGal-Leu plates supplemented with DMSO (0.02%) or with rapamycin at 100 or 200 nM. Plates were incubated at 30°C for 5 days.

**Supplementary Table S1.**
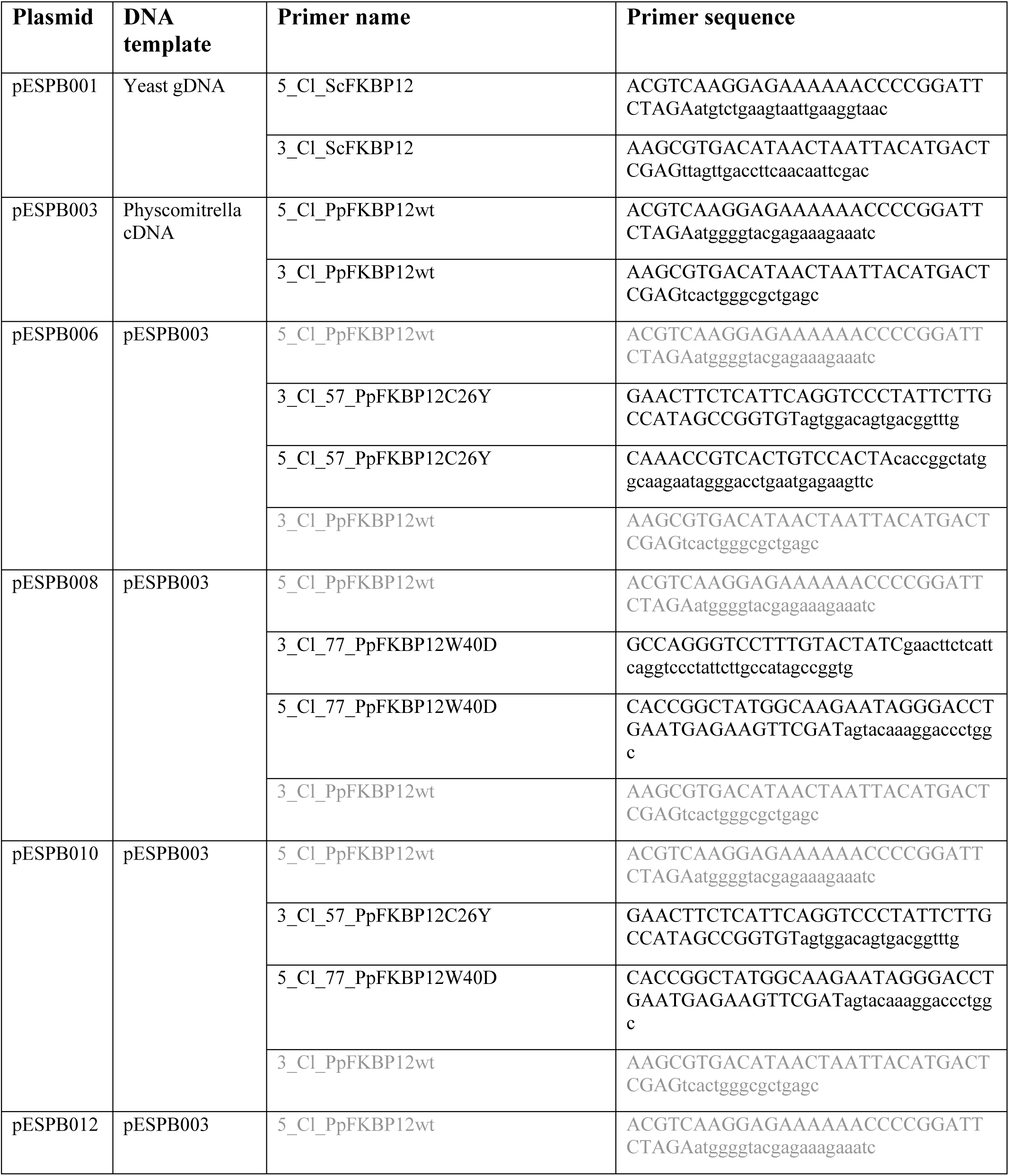
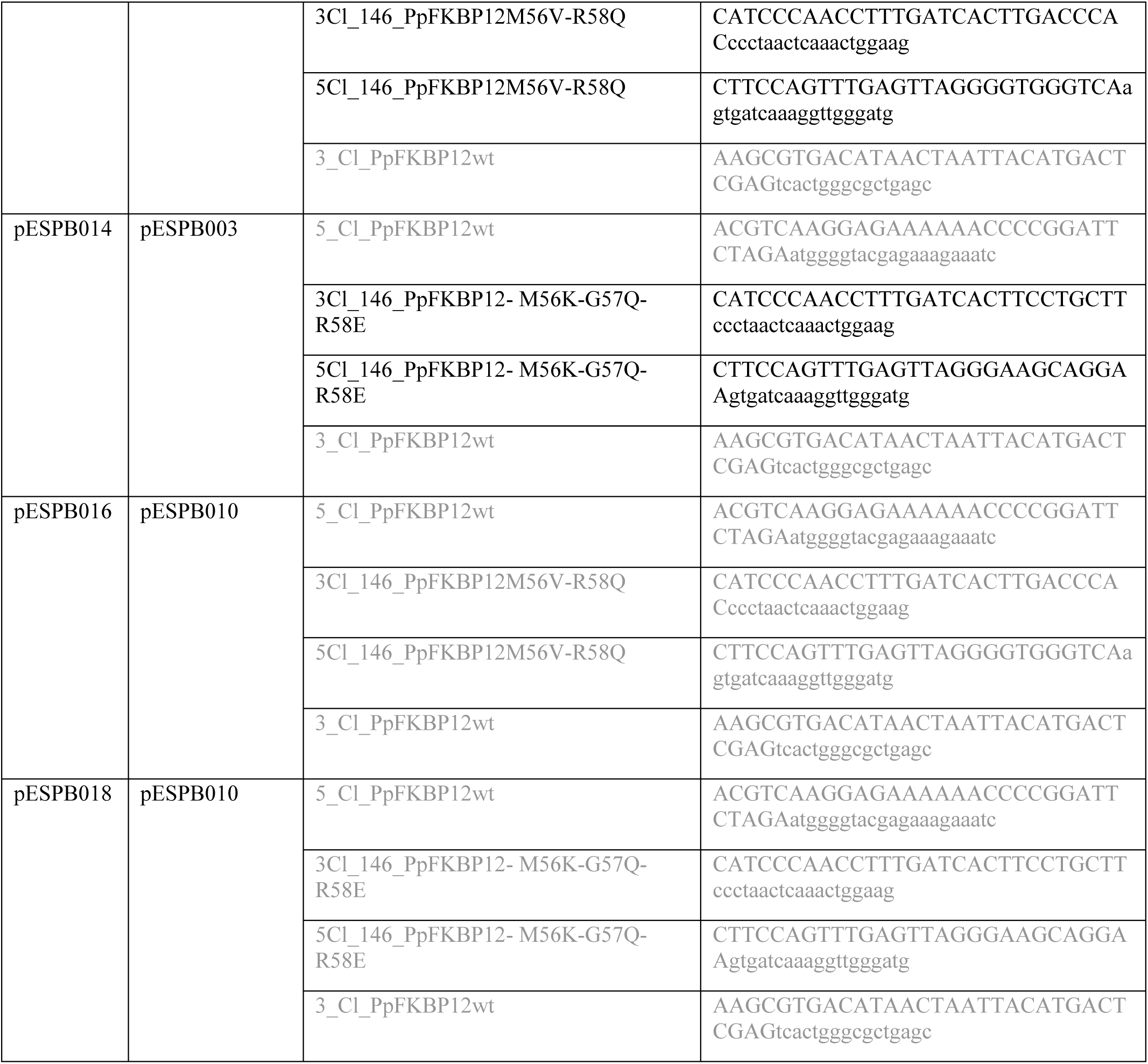
Primers used to clone *ScFKBP12*, *PpFKBP12* WT and mutants in p415 GAL1 restricted with XbaI and XhoI. Grey font colours indicate that the primers have been re-used.

**Supplementary Table S2.**
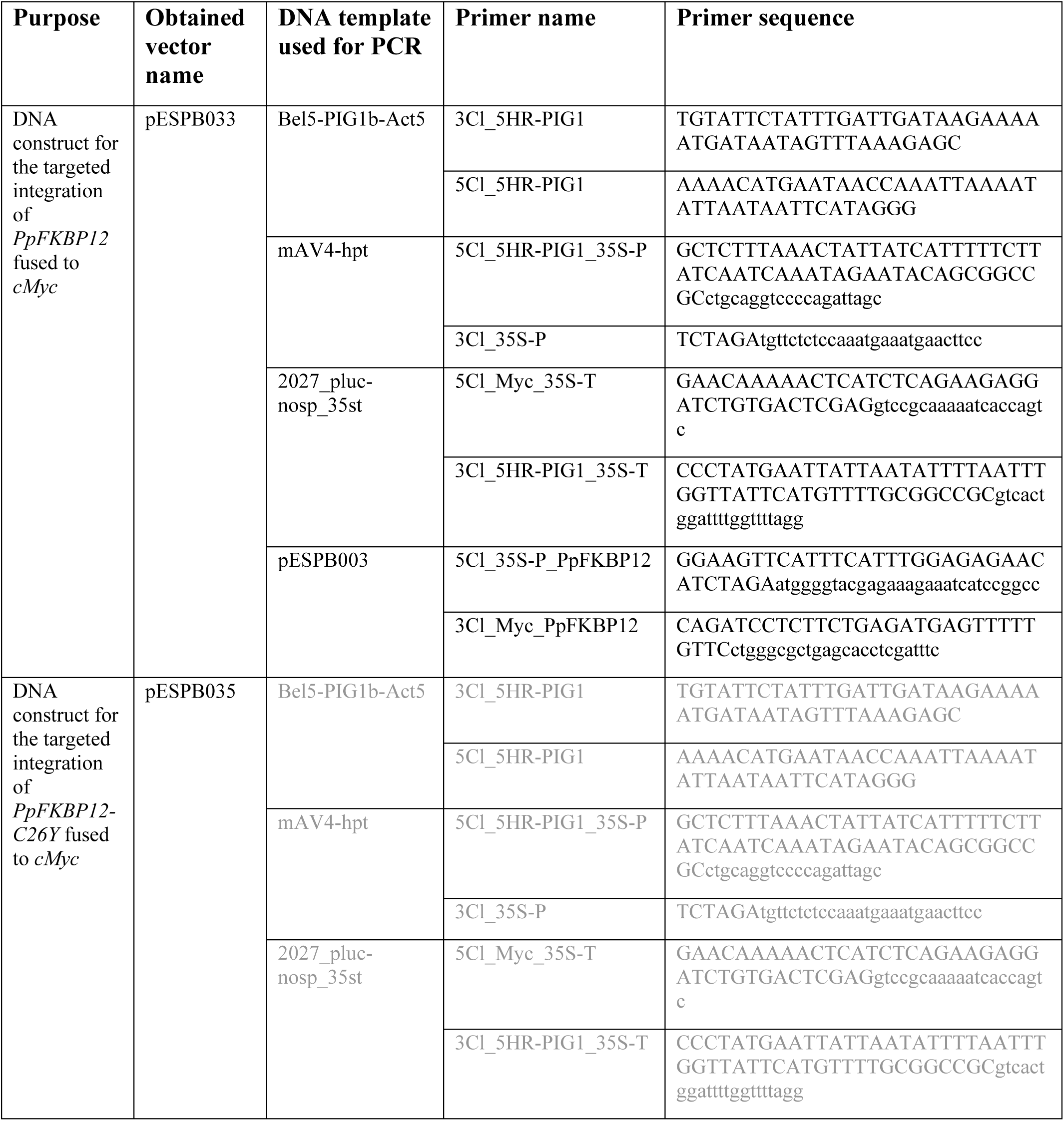
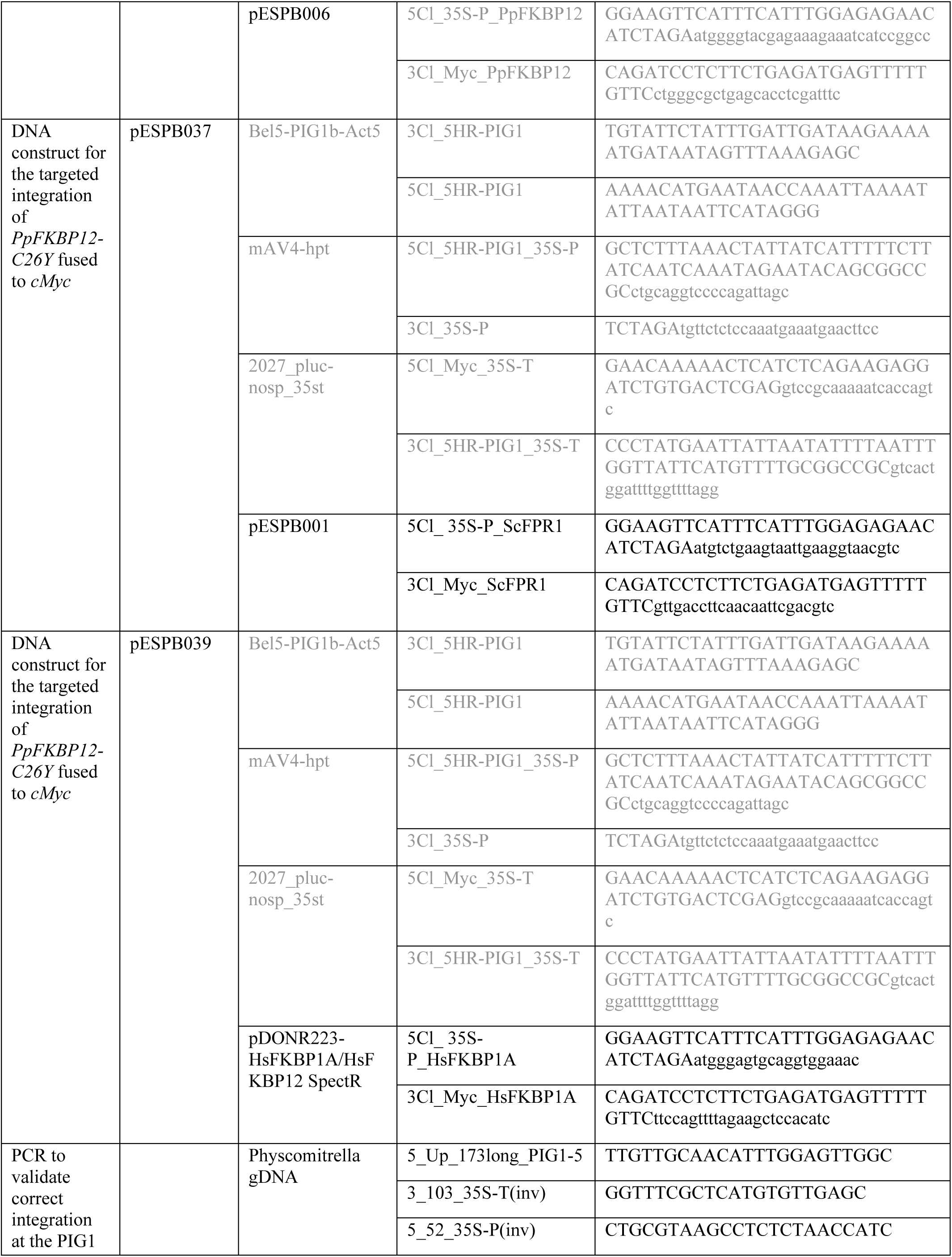
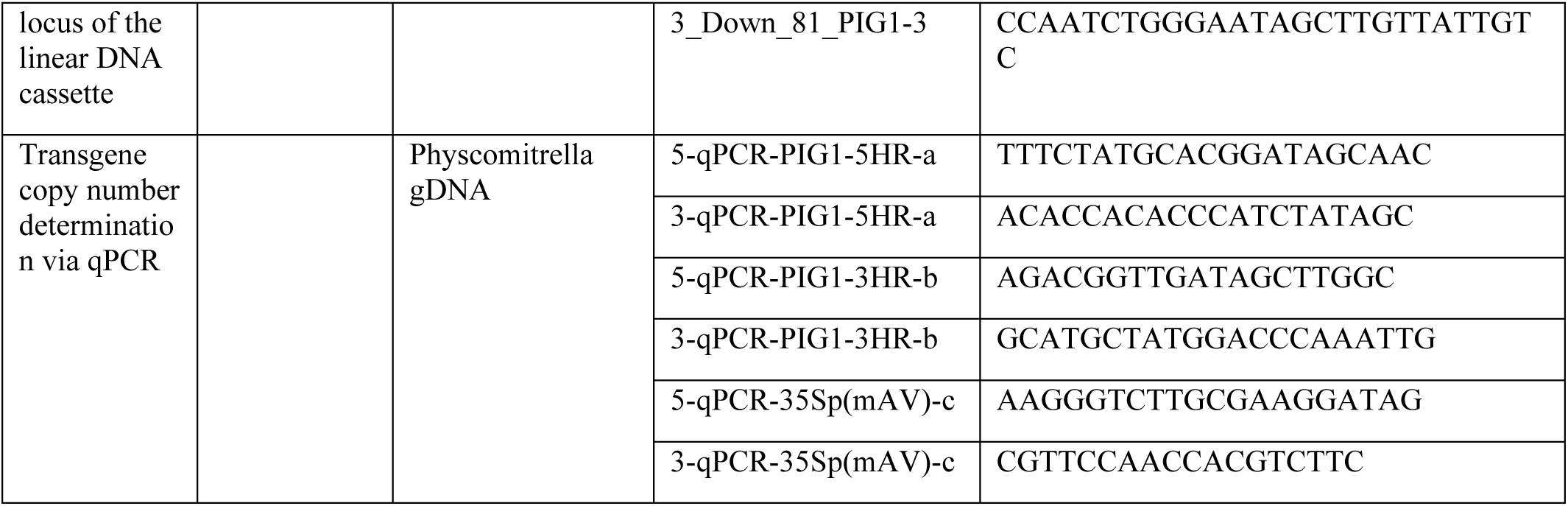
Primers used to generate the vectors containing the different *PIG1bL-CaMV 35Sp-FKBP12-cMyc-CaMV 35St-PIG1bR* DNA constructs, to validate via PCR their correct integration at the PIG1 locus, and to determine via qPCR their copy number in the Physcomitrella genome. Grey font colours indicate that the primers have been re-used.

**Supplementary Table S3.**
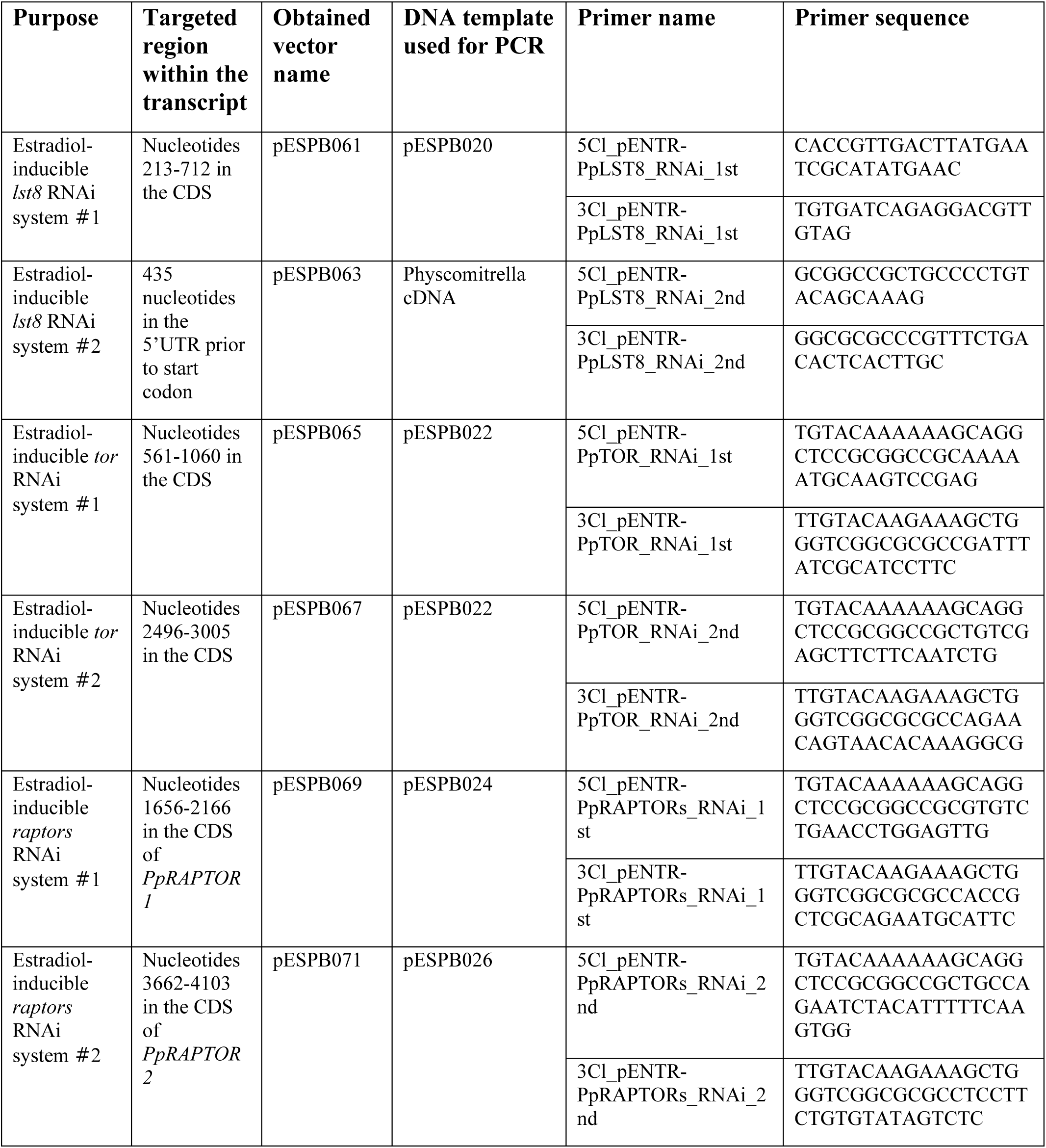
Primers used to generate the estradiol-inducible *lst8*, *tor*, and *raptors* RNA interference (RNAi) systems destined for a targeted integration at the PIG1 locus in the Physcomitrella genome.

**Supplementary Table S4.** International Moss Stock Center (IMSC, www.mossstock-center.org) accession numbers of the lines used in this study.

| Code | IMSC accession # | Genotype |
| --- | --- | --- |
| Wild type | 41269 |  |
| ESPB001 | 40976 | *PIG1bR-CaMV 35Sp-PpFKBP12-cMyc-CaMV 35St-PIG1bL* |
| ESPB002 | 40977 | *PIG1bR-CaMV 35Sp-PpFKBP12-cMyc-CaMV 35St-PIG1bL* |
| ESPB003 | 40978 | *PIG1bR-CaMV 35Sp-PpFKBP12-cMyc-CaMV 35St-PIG1bL* |
| ESPB004 | 40979 | *PIG1bR-CaMV 35Sp-PpFKBP12-cMyc-CaMV 35St-PIG1bL* |
| ESPB006 | 40980 | *PIG1bR-CaMV 35Sp-PpFKBP12<sup>C26Y</sup>-cMyc-CaMV 35St-PIG1bL* |
| ESPB007 | 40981 | *PIG1bR-CaMV 35Sp-PpFKBP12<sup>C26Y</sup>-cMyc-CaMV 35St-PIG1bL* |
| ESPB010 | 40982 | *PIG1bR-CaMV 35Sp-ScFKBP12-cMyc-CaMV 35St-PIG1bL* |
| ESPB011 | 40983 | *PIG1bR-CaMV 35Sp-ScFKBP12-cMyc-CaMV 35St-PIG1bL* |
| ESPB012 | 40984 | *PIG1bR-CaMV 35Sp-ScFKBP12-cMyc-CaMV 35St-PIG1bL* |
| ESPB014 | 40985 | *PIG1bR-CaMV 35Sp-HsFKBP12-cMyc-CaMV 35St-PIG1bL* |
| ESPB015 | 40986 | *PIG1bR-CaMV 35Sp-HsFKBP12-cMyc-CaMV 35St-PIG1bL* |
| ESPB016 | 40987 | *PIG1bR-CaMV 35Sp-HsFKBP12-cMyc-CaMV 35St-PIG1bL* |
| ESPB017 | 40988 | *PIG1bR-CaMV 35Sp-PpFKBP12-cMyc-CaMV 35St-PIG1bL* |
| ESPB018 | 40989 | *PIG1bR-CaMV 35Sp-PpFKBP12-cMyc-CaMV 35St-PIG1bL* |
| ESPB020 | 40990 | *Pplst8-RNAi-61'-40* |
| ESPB024 | 40991 | *Pplst8-RNAi-63'-38* |
| ESPB027 | 40992 | *Pptor-RNAi-65'-27* |
| ESPB032 | 40993 | *Pptor-RNAi-67'-17* |
| ESPB035 | 40994 | *Ppraptors-RNAi-69'-14* |
| ESPB040 | 40995 | *Ppraptors-RNAi-71'-25* |

